# Fatty acid scavenging enables cancer escape from KRAS inhibition

**DOI:** 10.64898/2026.04.01.715565

**Authors:** Zihang Yuan, Bo Lin, Chunlan Wang, Yingying Miao, Da Zhang, Ziru Meng, Gangqi Wang, Andrew M Lowy, Michael Karin, Fei Yang, Beicheng Sun, Hua Su

## Abstract

Although inhibitors of oncogenic KRAS have shown clinical efficacy^1^, resistance to KRAS inhibition is common^2^, and its molecular basis remains unclear. Here we show that KRASi-resistant cancer cells sustain mitochondrial bioenergetics through enhanced fatty acid (FA) metabolism, despite suppression of canonical KRAS signaling. Specifically, KRASi-resistant pancreatic cancer cells exploit macropinocytosis to scavenge FA released from adipose tissue, fueling beta-oxidation independently of KRAS-PI3Kα signaling. This adaptive metabolic program is driven by the adhesion G protein-coupled receptor ADGRB1, which activates non-canonical PI3Kγ-PAK1 signaling to stimulate macropinocytosis and maintain metabolic homeostasis under KRASi. Disruption of ADGRB1-PI3Kγ signaling dismantles this metabolic program and restores KRASi sensitivity. This pathway operates across multiple KRAS-mutated cancers and is associated with poor therapeutic response and outcome. These findings offer a promising strategy for overcoming KRASi resistance.

## Introduction

Pancreatic ductal adenocarcinoma (PDAC) remains one of the most lethal malignancies, with a five-year survival rate below 13% and few effective treatments^3^. The development of targeted PDAC interceptions has been hampered by genetic heterogeneity, dispersed oncogenic mutations, and relatively low disease incidence^4^. Nonetheless, recent therapeutic advances targeting ontogenically activated KRAS proteins, which are present in nearly 90% of PDACs^5^, where they enable metabolic reprogramming^6^, have shown considerable promise^7,8^. Structure-guided drug design had resulted in mutant-specific inhibitors such as sotorasib and adagrasib for KRAS^G12C^, which is common in lung cancer, and HRS-4642, MRTX1133 and RMC-9805 for KRAS^G12D^, a mutation found in ∼40% of patients with PDAC^1^. However, the clinical benefits of KRASi remain modest due to acquired drug resistance^9,10^. Until now the metabolic adaptations underlying KRASi resistance remain poorly understood, hindering the development of new approaches for restoring drug sensitivity and efficacy.

Macropinocytosis (MP) is an actin-driven endocytic process that allows cells engulf and utilize extracellular macromolecules as an energy source^11^, which is frequently upregulated in KRAS-driven cancers^12,13^. Here, we identify a hitherto unrecognized metabolic adaptation in which the adhesion G protein-coupled receptor ADGRB1/BAI1 engages the PI3Kγ-PAK1 axis to drive MP-dependent FA metabolism, thereby conferring KRASi resistance across multiple KRAS-mutant cancers.

### FA β-oxidation supports proliferation of cells resistant to KRAS^G12D^ inactivation

To define metabolic adaptions that enable resistance to KRASi, we first stratified human PDAC cell lines and patient-derived organoids harboring KRAS^G12D^ based on their response to the selective KRAS^G12D^ inhibitors HRS-4642 (HRS) and MRTX1133 (MRTX). We identified KRASi-sensitive (KS) PDAC cell lines and patient-derived organoids as well as a distinct subset of cell lines and organoids displaying intrinsic resistance (KR) (Extended Data Fig. 1a, b). Continuous drug exposure further generated acquired resistance (AR) models from initially sensitive cells and organoids (Extended Data Fig. 1c-e). Notably, both intrinsically and acquired KRASi resistance models showed effective suppression of canonical KRAS signaling, as indicated by reduced ERK and AKT phosphorylation following inhibitor treatment (Extended Data Fig. 1f-k). KR and AR cells continued to proliferate after genetic KRAS ablation (Extended Data Fig. 1l). These findings suggest that KRASi resistance is not driven by ineffective inhibition of KRAS signaling, instead reflecting adaptive mechanisms that sustain cell proliferation in the absence of KRAS function as a signaling protein.

To identify metabolic pathways selectively engaged by KR and AR cells, we performed RNA sequencing (RNA-seq) on KS AsPC-1 cells and KR PANC-1 *KRAS^G12D^*PDAC cells treated with KRASi. Comparative transcriptomic analysis revealed that KR cells adopt a distinct metabolic state, with lipid metabolism emerging as the most prominently altered pathway, followed by changes in carbohydrate, glycan, amino acid, and nucleotide metabolism (Extended Data Fig. 2a, b). Whereas KRASi suppressed expression of FA metabolic genes in KS cells, these genes were strongly upregulated in KR and AR cells (Extended Data Fig. 2c-e), suggesting that FA metabolism is the key adaptive mechanism supporting KRASi resistance. Because FA must first be converted into acyl-CoAs to enter downstream metabolic pathways^14^, we quantified acyl-CoA species via LC-MS/MS. KR and AR cells maintained elevated palmitoyl-CoA (C16:0) concentrations irrespective of KRAS signaling activity, in contrast to KS cells, in which palmitoyl-CoA levels were reduced by KRASi (Fig. 1a and Extended Data Fig. 2f). To determine whether exogenous FA fuel oxidative metabolism in AR cells, we performed stable isotope tracing using uniformly labeled ^13^C-palmitic acid (^13^C16:0-PA). AR cells showed markedly increased incorporation of labeled carbon into tricarboxylic acid (TCA) cycle intermediates—including citrate, α-ketoglutarate, succinate, and malate—compared with KS cells (Fig. 1b). KRAS inhibition suppressed PA-derived TCA flux in KS, but not KR or AR cells (Fig. 1b and Extended Data Fig. 2g). Consistent with sustained mitochondrial metabolism, KR/AR cells maintained elevated oxygen consumption (OCR), which was further augmented by the addition of PA, as well as increased mitochondrial ATP production, irrespective of KRAS inhibition (Fig. 1c, d and Extended Data Fig. 2h-k). These metabolic parameters were diminished by blocking CPT1-dependent FA mitochondrial import with the CPT1i etomoxir or CPT1 ablation, or by loss of the acyl-CoA synthetase ACSBG1, but remained unaffected by KRAS inhibition itself (Fig. 1c, d and Extended Data Fig. 2j-l). Conversely, KRASi robustly reduced oxidative metabolism in KS cells, indicating a fundamental divergence in metabolic dependency between the KR/AR and KS states (Fig. 1c, d and Extended Data Fig. 2h, i, k). At the cellular level, KR/AR cells and organoids exhibited increased mitochondrial content, lipid droplet (LD) accumulation, and enhanced mitochondrial-LD contacts—features that persisted under KRAS^G12D^ inhibition—whereas KS models showed the opposite pattern (Fig. 1e, f and Extended Data Fig. 3a, b). Mitochondrial gene expression followed a similar trend (Extended Data Fig. 3c). Together, these data demonstrate that KRASi-resistant PDAC cells can sustain mitochondrial energy production via enhanced FA oxidation.

**Fig. 1.**
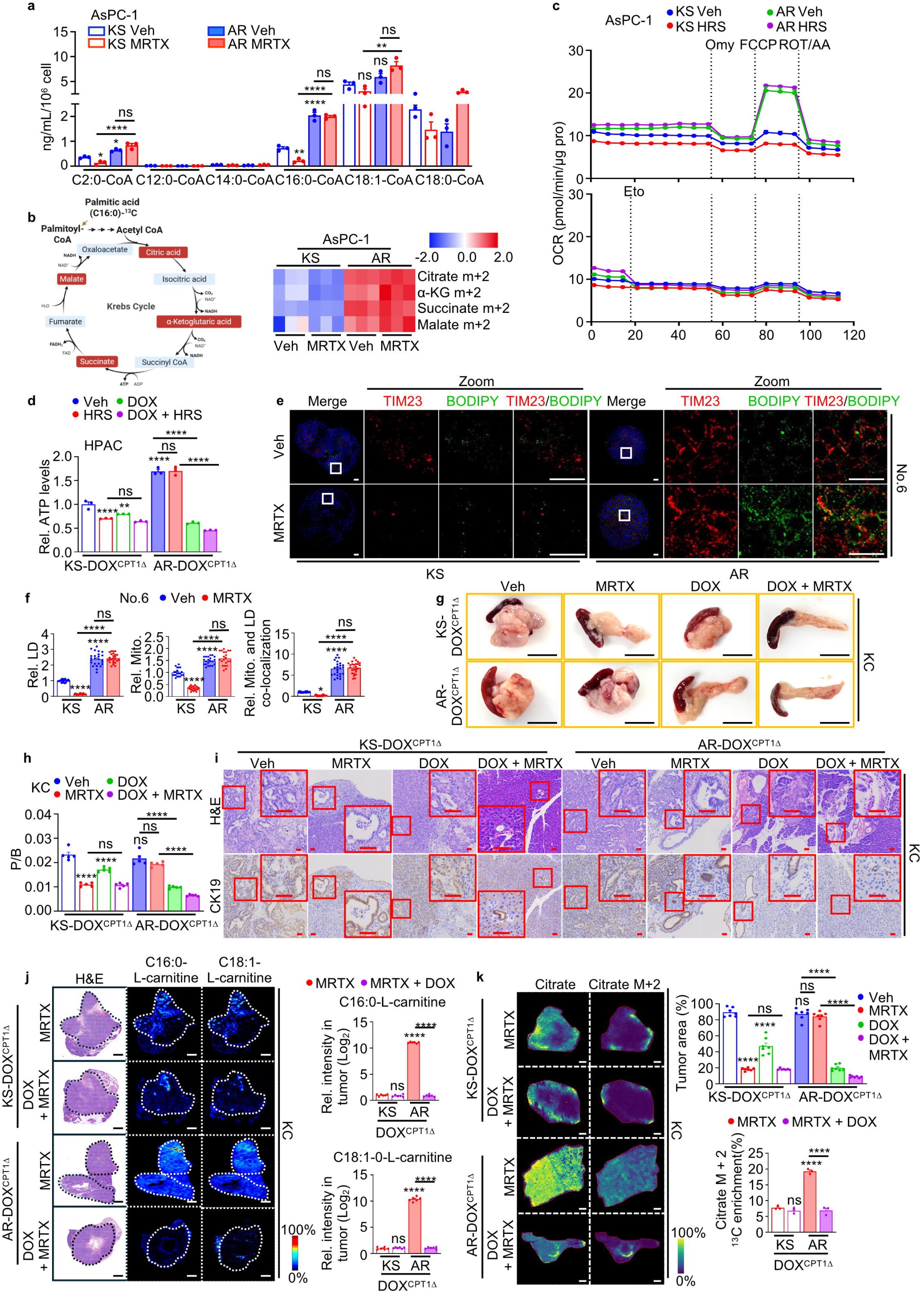
Fatty acid oxidation maintains the survival of KRASi-resistant PDAC cells. **a**, LC-MS analysis of intracellular acyl-coenzyme A species in sensitive (KS) and acquired (AR) AsPC-1 cells -/+ 100 nM MRTX1133 (MRTX) for 24 h. Veh, Vehicle. **b**, Right: Fractional labelling of TCA cycle intermediates in KS/AR AsPC-1 cells incubated with [U-^13^C]-PA and -/+ MRTX for 12 h. α-KG, α-ketoglutarate. Left: A schematic illustration of PA-derived acetyl-CoA fuels the TCA cycle. Blue, replicates with low expression; red, replicates with high expression. **c**, Oxygen consumption rate (OCR) of KS and AR AsPC-1 cells incubated -/+ PA and 100 nM HRS-4642 (HRS) for 24 h before and after addition of 40 μM Etomoxir (Eto), followed by oligomycin (Omy), FCCP, and rotenone/antimycin A (ROT/AA) treatments. **d**, KS/AR HPAC cells expressing doxycycline (DOX)-inducible CPT1 deletion (DOX^CPT1Δ^) were treated -/+ 0.2 μg/mL DOX for 48 h, and then -/+ HRS for 48 h. Total cellular ATP is relative to untreated KS cells. **e**, Representative images of mitochondria (TIM23) and BODIPY-labelled lipid droplets (LD) in parental KS No.6 and AR No.6 human PDAC organoids treated -/+ MRTX for 24 h. **f**, Quantification of mitochondria, LD, and their co-localization in (**e**). **g**, Pancreas morphology 3 weeks after orthotopic transplantation of KS and AR KC6141 (KC) cells expressing DOX^CPT1Δ^ and host treatment at 72 h post-transplantation -/+ 30 mg/kg MRTX (i.p.), 0.2 mg/mL DOX (drinking water), or DOX + MRTX. **h**, Pancreas weight relative to body weight (P/B weight) of mice in (**g**). **i**, Representative H&E and IHC staining for CK19 in pancreata from (**g**). Boxed areas are further magnified. Tumor areas are at the bottom. **j**, Representative AFADESI MSI images and quantification of endogenous C16:0 L-carnitine (m/z 400.3436) and C18:1 L-carnitine (m/z 426.3559) distribution in pancreata two weeks after orthotopic transplantation of KS/AR KC cells expressing DOX^CPT1Δ^. Host were treated starting 10 days post-transplantation with MRTX, DOX, or DOX + MRTX. AR tumors exhibit elevated FAO and increased sensitivity to CPT1 ablation compared to KS tumors. Boxed regions indicate tumor areas. **k**, Pancreata from mice in (**j**) were sectioned and cultured with 100 μM [U-^13^C]-PA for 4 h. Representative MALDI MSI images and quantification of unlabeled citrate (M + 0, m/z 191.019) and ^13^C-citrate (M + 2, m/z 193.026) distribution in pancreata, showing a similar trend to (**j**). Data in (**a**, **c**, **d**, **k**) (n=3 independent experiments), (**f**) (n=25 fields), (**h**) (n**=**5 mice), (**i**) (n=7 fields), and (**j**) (n=6 fields) are mean ± s.e.m. Statistical significance was determined using one-way analysis of variance (ANOVA) with Tukey post-hoc tests (**a**, **d**, **f**, **h**-**k**) based on data normality distribution. Exact *P* values are shown in the Source Data. \**P* < 0.05, \*\**P* < 0.01, \*\*\**P* < 0.001, \*\*\*\**P* < 0.0001. NS, not significant. Scale bars (**e**) 40 μm, (**g**) 1 cm, (**i**) 100 μm, (**j**, **k**) 500 μm.

To test whether FA oxidation (FAO) is required for KRASi-resistant tumor growth *in vivo*, we orthotopically transplanted KS or AR mouse KC6141 PDAC cells carrying a doxycycline (DOX)-inducible CPT1 ablation (CPT1^Δ^) cassette and treated the recipient mice with a KRASi and/or DOX. As expected, KRASi markedly suppressed KS tumor growth, evidenced by reduced pancreas-to-body weight ratios, diminished tumor area, and decreased expression of proliferation, FA-metabolism and mitochondrial-related markers (Fig. 1g-i, Extended Data Fig. 3d-f). In contrast, AR tumors showed opposite trends, remaining largely refractory to KRAS inhibition (Fig. 1g-i, Extended Data Fig. 3d-f). By contrast, CPT1 deletion impaired the growth of both KS and AR tumors, with a more pronounced effect in AR tumors (Fig. 1g-i, Extended Data Fig. 3d). Moreover, combined CPT1 deletion and KRASi treatment caused additional growth suppression, specifically in AR tumors (Fig. 1g-i), suggesting that inhibition of FAO restores KRASi sensitivity. Consistent with enhanced FAO, label-free, in situ AFADESI-MSI revealed elevated endogenous C16:0-L-carnitine and C18:1-L-carnitine, while MALDI-MSI detected increased citrate M+2 isotopologue abundance in AR tumors. These metabolic alterations were largely abolished by CPT1 ablation, demonstrating that FAO-derived acetyl-CoA fuels TCA cycle activity in AR tumors (Fig. 1j, k). Collectively, these results establish FAO as a critical metabolic dependency of KRASi-resistant PDAC.

### FA ingested via macropinocytosis enable metabolic adaption

To determine the source of FA sustaining the growth KR/AR cells, we first examined the contribution of *de novo* lipogenesis. Although deletion of FASN, a key enzyme in FA synthesis, suppressed the proliferation of both KS and AR cells, it did not sensitize either tumor type to KRASi (Extended Data Fig. 4a, b), indicating that endogenous FA synthesis is not the primary driver of KRASi resistance. Given the frequent presence of intrapancreatic adipose tissue in pancreatic disease and malignancy^15,16^, we next examined whether extracellular FAs support KR/AR survival. Disruption of major pancreatic FA transporter-associated proteins^17^ (CD36, FATP2, FATP3, and FABP5) selectively impaired KS cell proliferation, whereas culture in lipid-depleted medium selectively sensitized KR/AR cells to KRASi (Extended Data Fig. 4c-g). Compared to KS cells, AR cells also showed markedly increased uptake of fluorescently labeled PA (BODIPY-PA) (Extended Data Fig. 4h). These results suggest FA acquisition route in KR/AR cells independent of endogenous synthesis and FA transporters.

MP represents an alternative nutrient procurement pathway through which cancer cells take up extracellular nutrients, that plays a key role in PDAC metabolism and survival^11,12,18^. We therefore examined MP’s role in FA uptake by KRASi-resistant cells. AR PDAC cells exhibited markedly elevated MP rates, indicated by enhanced uptake of a high-molecular weight dextran (TMR-DEX) and its co-localization with BODIPY-PA, even during KRASi treatment. Pharmacological inhibition of MP with EIPA abrogated this uptake (Fig. 2a). In co-culture with adipocytes harboring BODIPY-PA, AR cells acquired more adipocyte-derived lipids than KS cells, irrespective of KRAS inhibition (Extended Data Fig. 4i). This effect was abolished by EIPA (Extended Data Fig. 4i), suggesting that AR cells scavenge FA released from surrounding cells in the microenvironment.

**Fig. 2.**
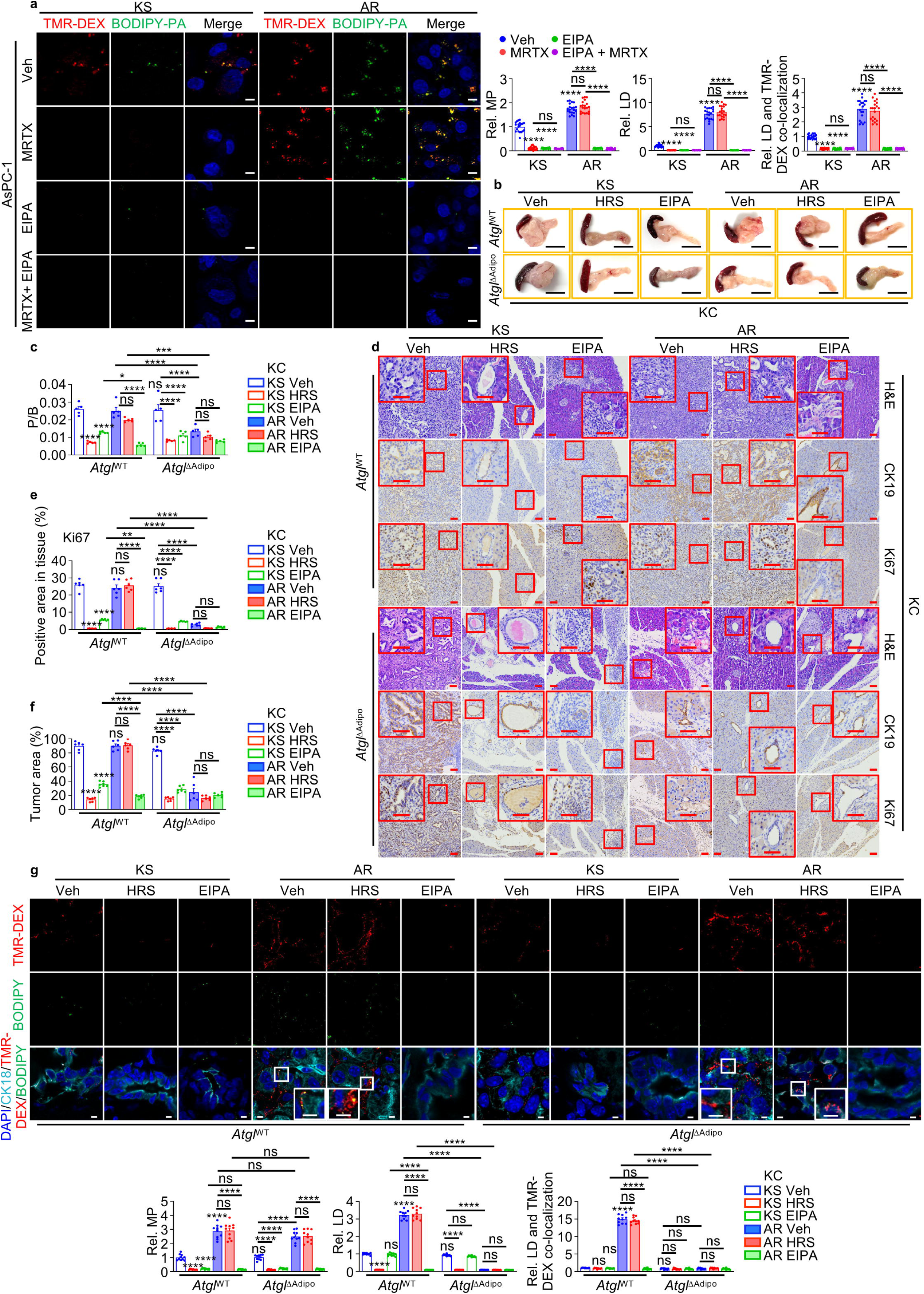
Macropinocytosis-mediated uptake of adipose-derived FA supports KRASi resistance. **a**, Representative images and quantification of BODIPY-labelled PA uptake, MP (TMR-DEX uptake), and their colocalization in KS and AR AsPC-1 cells -/+ MRTX, 25 μM EIPA, or MRTX + EIPA for 24 h. **b**, Pancreas morphology 3 weeks after orthotopic transplantation of KS and AR KC cells into *Atgl*^WT^ and *Atgl*^ΔAdipo^ mice -/+ 10 mg/kg HRS (i.p.) or 10 mg/kg EIPA (i.p.) 72 h post-transplantation. **c**, P/B weight ratio of mice from (**b**). **d**, Representative H&E and IHC of ductal (CK19) and proliferative (Ki67) markers in pancreata from (**b**). Boxed areas are further magnified. **e**, **f**, Image J quantification of Ki67 staining intensity (**e**) and tumor area (**f**) from (**d**). **g**, Representative images and quantification of MP, BODIPY-LD (BODIPY) and their co-localization in pancreata from (**b**). Pancreatic epithelial cells or carcinoma cells are marked by CK18 (cyan). Data in (**a**) (n=20 fields), (**c**, **g**) (n=5 mice), (**e**, **f**) (n=6 fields) and (**g**) (n=10 fields) are mean ± s.e.m. Statistical significance was determined using one-way ANOVA with Tukey post-hoc tests (**c**, **e**, **f**, and **g**) or Brown-Forsythe and Welch ANOVA tests with Dunnett T3 test (**a**, **g**) based on data normality distribution. Exact *P* values are shown in Source Data. \**P* < 0.05, \*\**P* < 0.01, \*\*\**P* < 0.001, \*\*\*\**P* < 0.0001. Scale bars (**a**) 10 μm, (**b**) 1 cm, (**d**) 100 μm, (**g**) 5 μm.

To directly test the requirement for adipocyte-derived FA in KRASi-resistant tumor growth, we orthotopically transplanted KS or AR cells into wild-type mice (*Atgl*^WT^) or mice specifically lacking ATGL (*Atgl*^ΔAdipo^), the lipase responsible for triglyceride hydrolysis^19^, in adipocytes. Loss of adipocyte ATGL markedly restrained AR tumor growth—reducing pancreas-to-body weight ratio, CK19 and Ki67 expression, and tumor area—independent of KRAS inhibition (Fig. 2b-f). Although MP activity remained high in AR tumors regardless of host genotypes or of KRAS inhibition, BODIPY-LD content and BODIPY-TMR-DEX colocalization were markedly reduced in transplanted KS and AR tumors from *Atgl*^ΔAdipo^ mice (Fig. 2g). Importantly, MP inhibition reduced tumor growth only in the presence of adipocyte-derived FA, exerting a profound inhibitory effect on AR tumors (Fig. 2b-f). Together, these findings suggest that MP-mediated uptake of adipocyte-derived FA sustains the growth of KRASi-resistant tumors and renders them KRAS independent.

### ADGRB1-PI3Kγ-pPAK1^S144^ signaling promotes FA uptake

Classical MP in PDAC cells is driven by KRAS-PI3Kα signaling^20^. However, the persistence of MP-mediated nutrient uptake by KRAS inhibited KR/AR cells prompted a search for alternative regulatory pathways. Transcriptomic profiling revealed upregulation of MP-associated genes, including *PIK3CG*, *CDC42*, *RAB5*, *RAB7*, and *SEPT6*, in AR cells, whereas expression of these genes was lower in KS cells and was barely affected by KRAS inhibition (Extended Data Fig. 4j). Notably, *PIK3CA* (encoding PI3Kα/p110α) expression was lower in AR than in KS cells and was further reduced by KRASi in both cell types. In contrast, *PIK3CG* (encoding PI3Kγ/p110γ)-a kinase implicated in lymphocyte MP and autophagy-deficiency activated NRF2-induced MP^12^-was substantially upregulated in AR cells and was not reduced by KRASi or NRF2 ablation (Extended Data Fig. 4j-l), suggesting PI3Kγ could be a mediator of KRAS- and NRF2-independent MP. Consistent with this notion, KRAS inhibition reduced phosphorylation of PAK1 at T423, a PI3Kα-dependent site^21^, in both AR and KS cells, whereas PI3Kγ-dependent pS144-PAK1^22^ was elevated only in AR cells and unaffected by KRAS inhibition, indicating a KRAS-independent PI3Kγ-PAK1 axis (Extended Data Fig. 4k). Moreover, inhibition of PI3Kα with LY294002 and KRASi selectively suppressed adipocyte-derived lipid uptake and FA β-oxidation in KS cells, whereas the PI3Kγ inhibitor IPI549 selectively impaired them in AR cells (Extended Data Fig. 4m, n). This suggests that KRASi-resistant cells engage non-canonical PI3Kγ-PAK1 signaling to sustain KRAS independent MP and lipid metabolism.

To investigate how PI3Kγ-PAK1 signaling is activated in KR/AR cells, we re-examined our RNA-seq data and found significant enrichment of G protein-coupled receptor (GPCR)-G protein signaling in KR cells. This included upregulation of four GPCRs, ADRA2C, GPRC5B, ADGRB1, and GPR158, together with multiple G proteins (Extended Data Fig. 5a), that are normally activated by ligand-bound GPCRs^23^. Systematic GPCR ablation revealed that only loss of ADGRB1, an adhesion GPCR with largely undefined pathophysiology^24^, selectively reduced pS144-PAK1, without affecting pT423-PAK1 (Fig. 3a). ADGRB1 expression was upregulated across multiple AR PDAC models relative to KS models (Extended Data Fig. 5b, c). ADGRB1 engages downstream signaling by recruiting Gα12/13 and Gβγ subunits^25^. Using a Gα12/13 activity reporter (SRF-RE luciferase)^25^ we observed a marked increase in Gα12/13 activity in AR cells, which was abolished on ADGRB1 ablation (Extended Data Fig. 5d, e), an effect not seen in KS cells. *In vivo*, AR tumors exhibited enhanced uptake of adipocyte-derived lipids compared to KS tumors, irrespective of KRAS inhibition and this enhanced uptake was reversed by loss of ADGRB1 (Fig. 3b). In addition, ADGRB1 ablation selectively reduced MP, and suppressed pS144-PAK1 in AR cells (Fig. 3c, d). These phenotypes were rescued by expression of a constitutively active PI3Kγ^R1021C^ variant^26^, indicating that MP activation in KRASi-resistant cells depends on ADGRB1-PI3Kγ-pS144-PAK1 signaling (Fig. 3c, d). Consistently, ^13^C-PA tracing and OCR analysis showed that ADGRB1 loss impaired FAO selectively in AR cells (Fig. 3e, Extended Data Fig. 5f), an effect that was reversed by PI3Kγ^R1021C^ expression and blunted by EIPA treatment (Fig. 3e). These results identify ADGRB1 as an upstream activator of KRAS-independent PAK1 signaling that sustains MP-mediated-FA metabolism in KRASi resistant tumors.

**Fig. 3.**
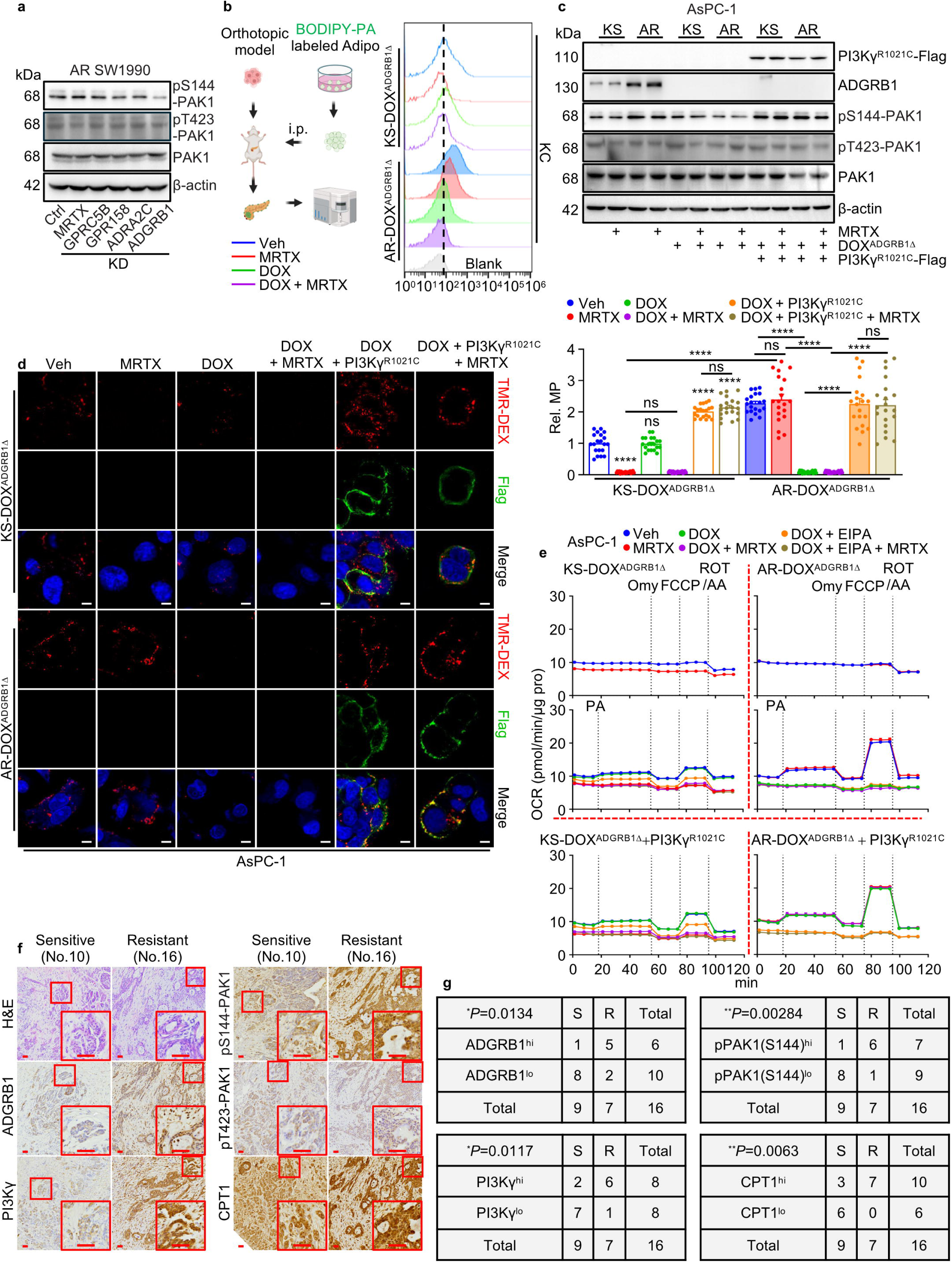
ADGRB1-PI3Kγ-pS144-PAK1 driven MP fuels KRASi resistance. **a**, Immunoblot (IB) analysis of indicated proteins in acquired KRASi-resistant SW1990 cells (AR) -/+ knock down (KD) of indicated GPCRs and -/+ MRTX for 2 h. **b**, AR and KS KC cells expressing DOX^ADGRB1Δ^ were orthotopically transplanted into mice -/+ 7 days MRTX, DOX, or DOX + MRTX, initiated 72 h post-transplantation. Then the BODIPY signal in PDAC cells was analyzed by flow cytometry 24 h after i.p. of BODIPY-PA loaded adipocytes (3T3-L1). The experimental scheme is shown to the left. **c**, IB analysis of indicated proteins in the indicated KS and AR AsPC-1 cells -/+ Flag-tagged PI3Kγ^R1021C^ expression and -/+ DOX treatment for 48 h, followed by -/+ MRTX for 2 h. **d**, Representative images and quantification of MP in TMR-DEX-incubated cells from (**c**). **e**, OCR of cells in (**c**) treated -/+ DOX for 48 h, followed by -/+ MRTX or EIPA for 24 h before and after -/+ PA incubation, and treatment with Omy, FCCP, and rotenone/antimycin A. **f**, Representative H&E and IHC staining of resected human PDAC tissues classified as KS or KR based on organoid sensitivity to KRASi (Extended Data Fig. 1b). Boxed areas are further magnified. **g**, Correlations between the indicated proteins and KRASi sensitivity in (**f**) were analyzed by a two-tailed Chi-square test. Data in **b** (n=3 mice), **d** (n=20 fields) and **e** (n=3 independent experiments) are mean ± s.e.m. Statistical significance was determined using Brown-Forsythe and Welch ANOVA tests with Dunnett T3 test (**d**) based on data normality distribution. \**P* < 0.05, \*\**P* < 0.01, \*\*\**P* < 0.001, \*\*\*\**P* < 0.0001. Exact *P* values are shown in Source Data. Scale bars (**d**) 10 μm, (**g**) 100 μm.

### ADGRB1 correlates with KRASi resistance and poor clinical outcome

To assess the clinical relevance of the ADGRB-PI3Kγ-PAK1 signaling axis, we examined its association with KRAS inhibitor response and patient outcomes in human pancreatic cancer. Immunohistochemistry (IHC) analysis of surgically resected human PDAC with KRAS^G12D^ mutation (the corresponding organoids shown in Extended Data Fig.1b) showed that most intrinsic KR tumors exhibited stronger staining for ADGRB1 (5/7), PI3Kγ (6/7), pPAK1 (S144) (6/7) and CPT1 (7/7) than did KS tumors (Fig. 3f, g), suggesting patients with high expression of these proteins were more likely to become KRASi resistant. In addition, ADGRB1 and pS144-PAK1 showed a significant positive correlation (Extended Data Fig. 5g-i). This relationship was independently validated in an additional cohort of 42 patients with PDAC, despite the absence of information regarding their KRASi responsiveness. Importantly, patients whose tumors were enriched for ADGRB1 or pS144-PAK1 had poorer median survival than patients with low ADGRB1 or pS144-PAK1 expression (Extended Data Fig. 5j). These results are consistent with the findings made in our preclinical models, suggesting that ADGRB1-activated signaling and FA metabolism may also drive KRASi resistance in human PDAC.

### The ADGRB1-PI3Kγ axis mediates general KRASi resistance

Oncogenic KRAS mutations are common across multiple cancer types^1^. We asked whether activation of the ADGRB1-PAK1 axis represents a generalizable mechanism of KRASi resistance beyond the KRAS^G12D^ mutation and PDAC. To address this, we examined additional KRAS inhibitor-resistant cancer models spanning distinct KRAS alleles and tissue origins. We first evaluated KRAS^G12C^-mutant PDAC cells (MIA-PaCa-2), and KRAS^G12D^-mutant human endometrial adenocarcinoma (HEC-1-B) and colon adenocarcinoma (GP2D) cells, stratifying each model into KRASi-sensitive (KS) or KRASi-resistant (KR/AR) (Extended Data Fig. 6a). Across all resistant models, AR MIA-PaCa-2, AR GP2D, and HRS-treated KR HEC-1-B cells displayed pronounced upregulation of FA metabolic genes, activation of ADGRB1-PI3Kγ-pS144-PAK1 signaling, and elevated Gα12/13 activity relative to KS counterparts or untreated KR cells (Fig. 4a, Extended Data Fig. 6b, c). ADGRB1 deletion reduced Gα12/13 activity only in AR cells (Extended Data Fig. 6b). Accordingly, AR MIA-PaCa-2, GP2D, and HRS-treated KR HEC-1-B cells displayed higher MP rates, FAO, and ATP production than KS or untreated cells (Fig. 4b-e, Extended Data Fig. 6d-j). These phenotypes were largely reversed in KR/AR cells by deletion of ADGRB1 or inhibition of PI3Kγ, whereas inhibition of KRAS or PI3Kα suppressed these parameters only in KS cells (Fig. 4b-e, Extended Data Fig. 6d-j) indicating convergence across tissues and cancers types.

**Fig. 4.**
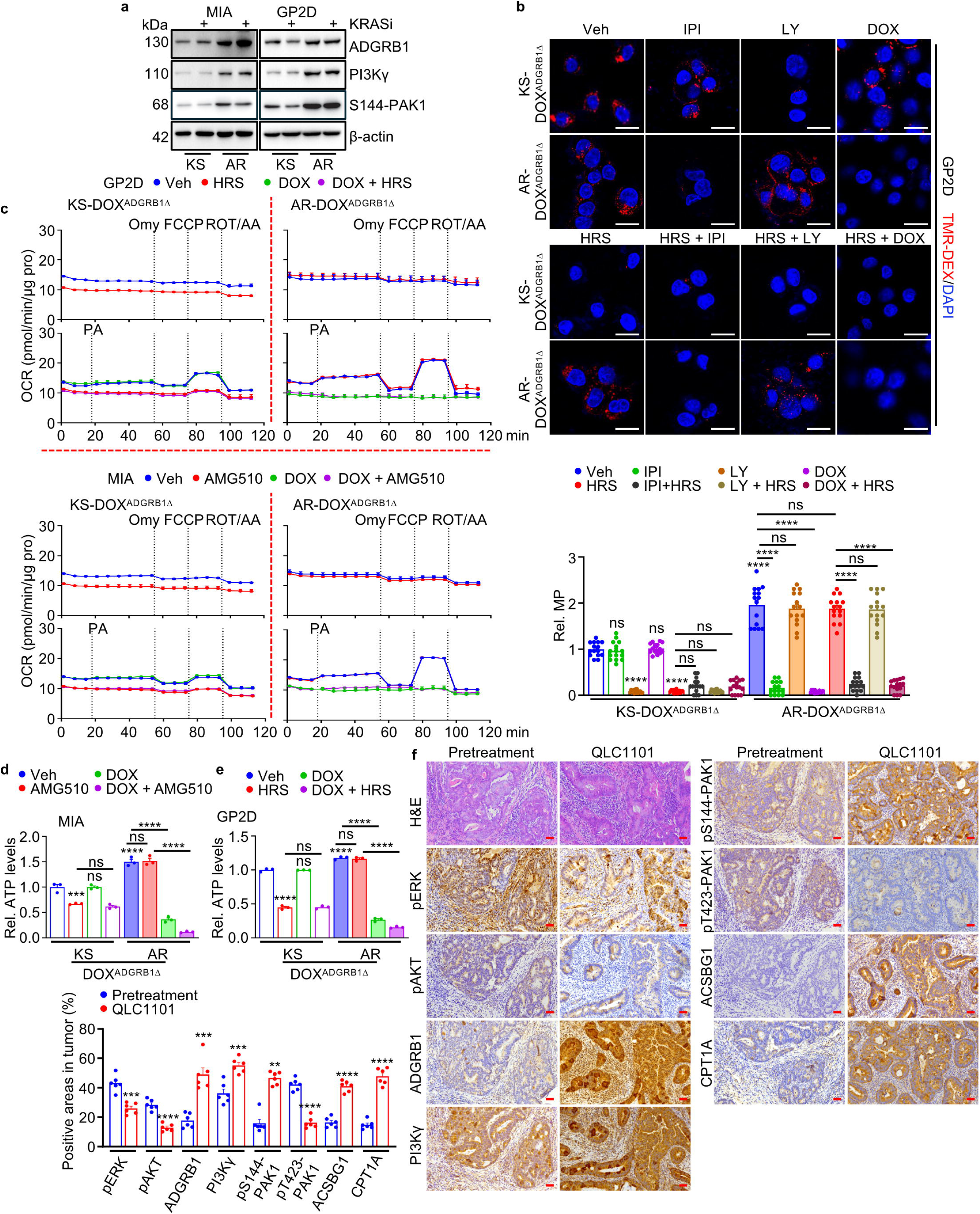
ADGRB1 signaling sustains energy production across diverse KRASi-resistant cancers. **a**, IB analysis of indicated proteins in KRAS^G12C^ mutated KS and AR MIA PaCa-2 cells (MIA) or KRAS^G12D^ mutated GP2D cells -/+ 100 nM of the KRAS^G12C^i AMG510 or the KRAS^G12D^i HRS for 24 h. **b**, Representative images and quantification of MP in KS/AR GP2D cells expressing DOX^ADGRB1Δ^ treated -/+ DOX for 48h, followed by treatment -/+ 1 μM PI3Kγi IPI549 (IPI), 1 μM PI3Kαi LY494002 (LY), HRS, HRS + IPI, or HRS + LY for 24 h. **c**, OCR of indicated cells -/+ DOX for 48h, followed by -/+ AMG510 (MIA) or HRS (GP2D) for 24 h, before and after -/+ PA, followed by Omy, FCCP, and rotenone/antimycin A treatments. **d**, **e**, Total cellular ATP in KS and AR MIA PaCa-2 or GP2D cells expressing DOX^ADGRB1Δ^ treated with the indicated compounds for 48h. **f**, Representative IHC and quantification in resected human colon adenocarcinoma tissues before and after KRAS^G12D^i QLC1101 treatment. Data in (**b**) (n=15 fields), (**c**, **d**, **e**) (n=3 independent experiments), and (**f**) (n=6 fields) are mean ± s.e.m. Statistical significance was determined using one-way ANOVA with Brown-Forsythe and Welch corrections followed by Dunnett T3 test (**b**), or with Tukey post-hoc tests (**d**, **e**), or two-sided unpaired *t*-test or Mann-Whitney *U*-test (**f**) based on data normality distribution. \**P* < 0.05, \*\**P* < 0.01, \*\*\**P* < 0.001, \*\*\*\**P* < 0.0001. Exact *P* values are shown in Source Data. Scale bars (**b**) 20 μm, (**f**) 100 μm.

For proof of concept as to whether this pathway is engaged in human tumors with acquired clinical resistance, we analyzed tumor specimens from a patient with KRAS^G12D^-mutant colorectal cancer who developed resistance after prolonged treatment with the KRAS^G12D^ inhibitor QLC1101. Compared with the corresponding pre-treatment tumor, the QLC1101-resistant tumor exhibited reduced levels of pERK, pAKT, and pT423-PAK1 confirming suppression of canonical KRAS signaling. In contrast, expression of ADGRB1-pS144-PAK1 signaling components and the FA metabolic enzymes CPT1A and ACSBG1 were markedly elevated (Fig. 4f). Together, these findings suggest that ADGRB1 signaling represents a common resistance mechanism co-opted by distinct KRAS-mutated cancers.

### ADGRB1-PI3Kγ-pS144-PAK1 targeting restores KRASi sensitivity

Given that ADGRB1-driven PI3Kγ signaling sustains MP and FAO in KR cells, we asked whether disrupting the ADGRB1-PI3Kγ-pS144-PAK1 pathway can restore KRASi sensitivity. To address this, we genetically or pharmacologically inhibited components of this signaling pathway across multiple KR/AR and KS cancer models. Deletion of ADGRB1 or CPT1, or PI3Kγ inhibition resensitized multiple KR/AR cancer cell models to KRASi, whereas these perturbations had minimal effects in KS cells, irrespective of KRASi treatment (Extended Data Fig. 6k-m). Combined PI3Kγ and KRAS inhibition produced a pronounced synergistic effect in AR models. SynergyFinder^+27^ analysis yielded median Bliss scores of 14.62 (AR No.6) and 12.43 (AR AsPC-1), in contrast to negligible scores in matched KS cells (0.26 in KS No.6; 0.93 in KS AsPC-1), indicating a synergy (score >10) restricted to resistant states (Fig. 5a, Extended Data Fig. 7a). This synergy was most prominent in resistant models treated with 10-1000 nM IPI and 10-10,000 nM HRS, whereas sensitive models which were highly responsive to HRS monotherapy displayed no meaningful synergy across the tested dose range (Fig. 5a, Extended Data Fig. 7a). Consistently, ADGRB1 deletion selectively suppressed AR organoid growth and triggered profound organoid regression when combined with KRASi (Extended Data Fig. 7b).

**Fig. 5.**
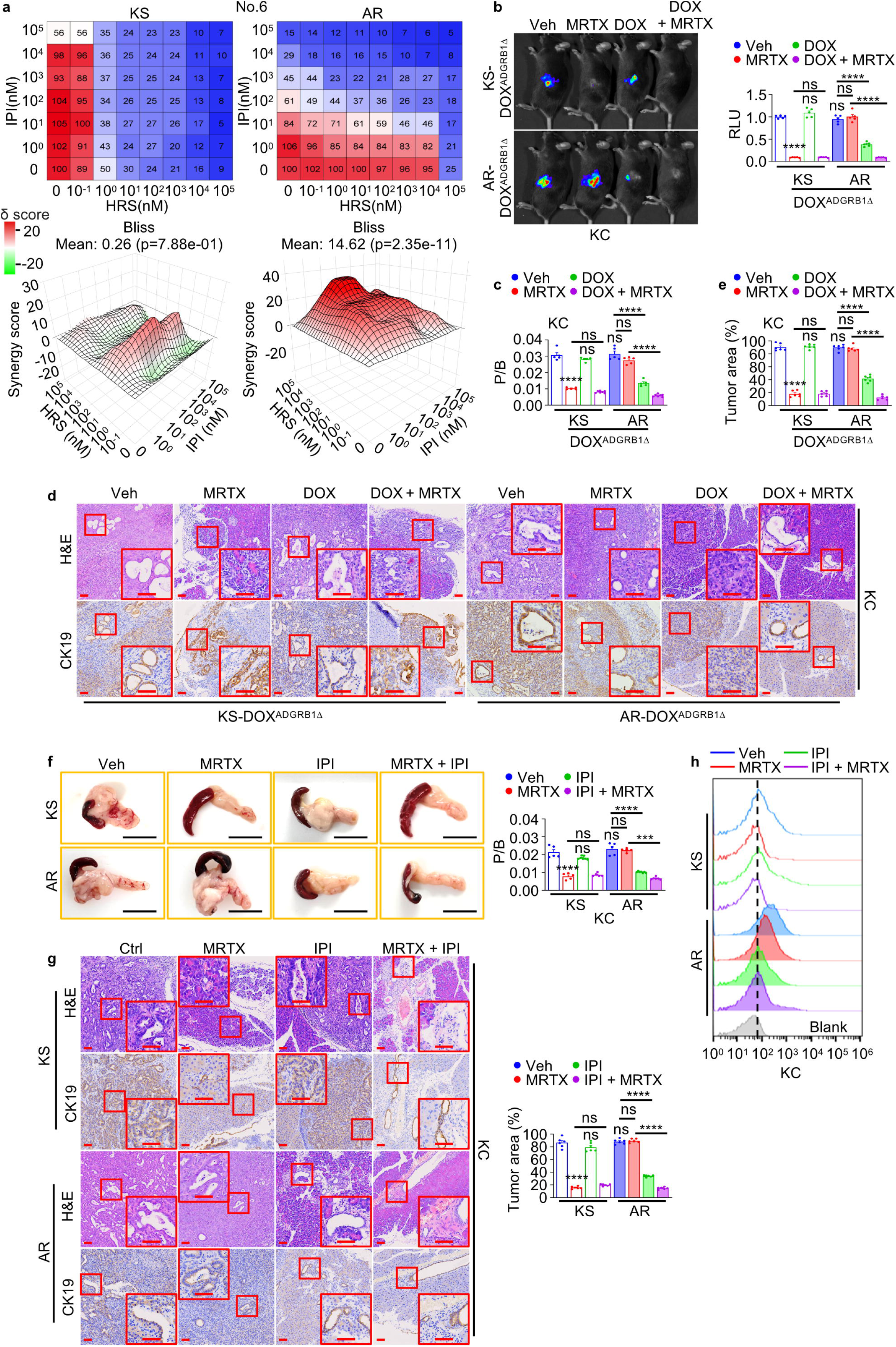
Inhibition of ADGRB1 signaling induces regression of KRASi-resistant tumors. **a**, KS/AR No.6 organoids were treated with IPI and/or HRS at the indicated concentrations for 7 days, and cell viability was assessed with CellTiter-Glo (CTG). Upper: Dose-response matrix (viability) for IPI and HRS. Blue intensity indicates the degree of inhibition. Lower: 3D synergy plots generated by SynergyFinder^+^. The synergy score is shown by red (>0) and green (<0). The synergy score > 10 indicates strong synergic effect. **b**, Akaluciferase bioluminescence imaging of mice 3 weeks after orthotopic transplantation of KS or AR KC cells expressing DOX^ADGRB1Δ^ and a Ki67-akaluciferase reporter, -/+ MRTX, DOX, or DOX + MRTX treatments initiated 72 h post-transplantation. Quantification of akaluciferase activity, expressed as relative light units (RLU), is on the right. **c**, P/B ratio of mice from (**b**). **d**, Representative H&E and CK19 IHC staining of pancreata from (**b**). Boxed areas are further magnified. **e**, Quantification of tumor areas from (**d**). **f**, Pancreas morphology and P/B ratio 3 weeks after orthotopic transplantation of indicated cells into mice -/+ MRTX, 15 mg/kg IPI (p.o.), or MRTX + IPI treatments initiated 72 h post-transplantation. **g**, Representative H&E and CK19 staining of pancreata from (**f**). Boxed areas are further magnified. Quantification of tumor areas is shown to the right. **h**, Flow cytometry analysis of BODIPY signals in PDAC cells harvested 24 h after i.p. injection of BODIPY-PA-loaded adipocytes into mice from (**f**) at day 7 post-KS and AR KC cell implantation. Data in (**b**) (n=5 mice), (**c**, **f**) (n=5-7 mice), (**e**, **g**) (n=6 fields), and (**h**) (n=3 mice) are mean ± s.e.m. Statistical significance was determined using one-way ANOVA with Tukey post-hoc tests. \**P* < 0.05, \*\**P* < 0.01, \*\*\**P* < 0.001, \*\*\*\**P* < 0.0001. Exact *P* values are shown in Source Data. Scale bars (**d**, **g**) 100 μm, (**f**) 1 cm.

To probe the contribution of ADGRB1-PI3Kγ-pS144-PAK1 signaling to *in vivo* AR tumor growth, KS or AR cells harboring a DOX-inducible ADGRB1 ablation cassette and a Ki67-driven Aka-luciferase reporter^28^ were orthotopically transplanted into mice that were either treated with MRTX and/or DOX. Whereas MRTX suppressed only KS tumors, ADGRB1 depletion or PI3Kγ inhibition selectively impeded AR tumor growth, as reflected by reduced luciferase activity, decreased pancreas-to-body weight ratio, diminished CK19 staining and tumor areas, and was accompanied by lower pS144-PAK1 expression (Fig. 5b-g, Extended Data Fig. 7c, d). Combined inhibition of ADGRB1 or PI3Kγ with KRAS further triggered AR tumor regression (Fig. 5b-g). Accordingly, lipid uptake was elevated in AR relative to KS tumors, irrespective of KRAS inhibition, and this was reversed by PI3Kγ inhibition (Fig. 5h). Together, these findings demonstrate that ADGRB1-PI3Kγ signaling is both necessary and therapeutically actionable for maintaining KRASi resistance.

## Discussion

Although cancer cells are known to develop KRASi resistance through multiple mechanisms, including *de novo* KRAS synthesis^29^, activation of IRE1α-dependent proteostasis^30^, enhanced proteasomal degradation^31^, and reactivation of upstream signaling via EGFR^32^, it is indisputable that KRAS activity can still be suppressed by KRASi. How cancer cells maintain metabolic homeostasis while overcoming KRAS inhibition was previously unknown. We now show that activation of ADGRB1-stimulated FA metabolism is a general mechanism of resistance to KRASi across distinct KRAS mutations and cancer types. Although elevated tissue and circulating lipid levels were documented in patients with PDAC^33,34^, the contribution of MP-mediated scavenging of adipocyte-derived FA to fuel the survival and growth of KRASi-resistant cells was previously unknown. Adipocyte-derived FA may also serve as a lipid reservoir for other components of the tumor microenvironment, such as cancer-associated fibroblasts^35^. Beyond lipids, MP can also internalize other macromolecules, such as collagens^18^, thereby supporting the metabolic demands of KR/AR cells. Indeed, our transcriptomic analyses reveal coordinated upregulation of amino acid and nucleotide metabolic pathways (Extended Data Fig. 2b), indicative of a broad and flexible metabolic rewiring that enables KR/AR tumor cell survival and growth. Although PI3Kγ has been implicated in activation of MP in lymphocytes, downstream of NRF2 signaling in autophagy-deficient cancer cells^12^ and in KRAS^G12R^-mutant contexts^36^, its role in MP in KR/AR cells was not previously explored. Notably, PI3Kγ expression in KRASi-resistant cells appears to be independent of NRF2 (Extended Data Fig. 4l), suggesting a distinct regulatory mechanism in this setting. Our results uncover a previously unappreciated role for the ADGRB1-PI3Kγ-pS144-PAK1 signaling axis in sustaining MP in the absence of KRAS-PI3Kα signaling. They also raise the possibility that PI3Kγ inhibition by IPI549 represents an immediately tractable strategy to test in the clinic. In addition, these findings highlight a key role for MP in mediating interactions between cancer cells and the tumor microenvironment under diverse tumor stress conditions.

ADGRB1 is an underexplored adhesion GPCR, previously studied as a phagocytic receptor in macrophages that recognizes phosphatidylserine on apoptotic cells^37^. ADGRB1’s role in the activation of MP and FA metabolism in epithelial cancers substantially expands the functional repertoire of adhesion GPCRs^38^. The coordinated upregulation of ADGRB1, lipid metabolism-associated, and mitochondrial genes during the emergence of KRASi resistance in the absence of detectable gain-of-function mutations by whole-exome sequencing, could be explained by a yet-to-identified epigenetic or transcriptional regulatory mechanism. Another important challenge is the identification of ADGRB1 antagonists as this will enable development of compounds capable of reversing KRASi resistance.

**Table 1:**
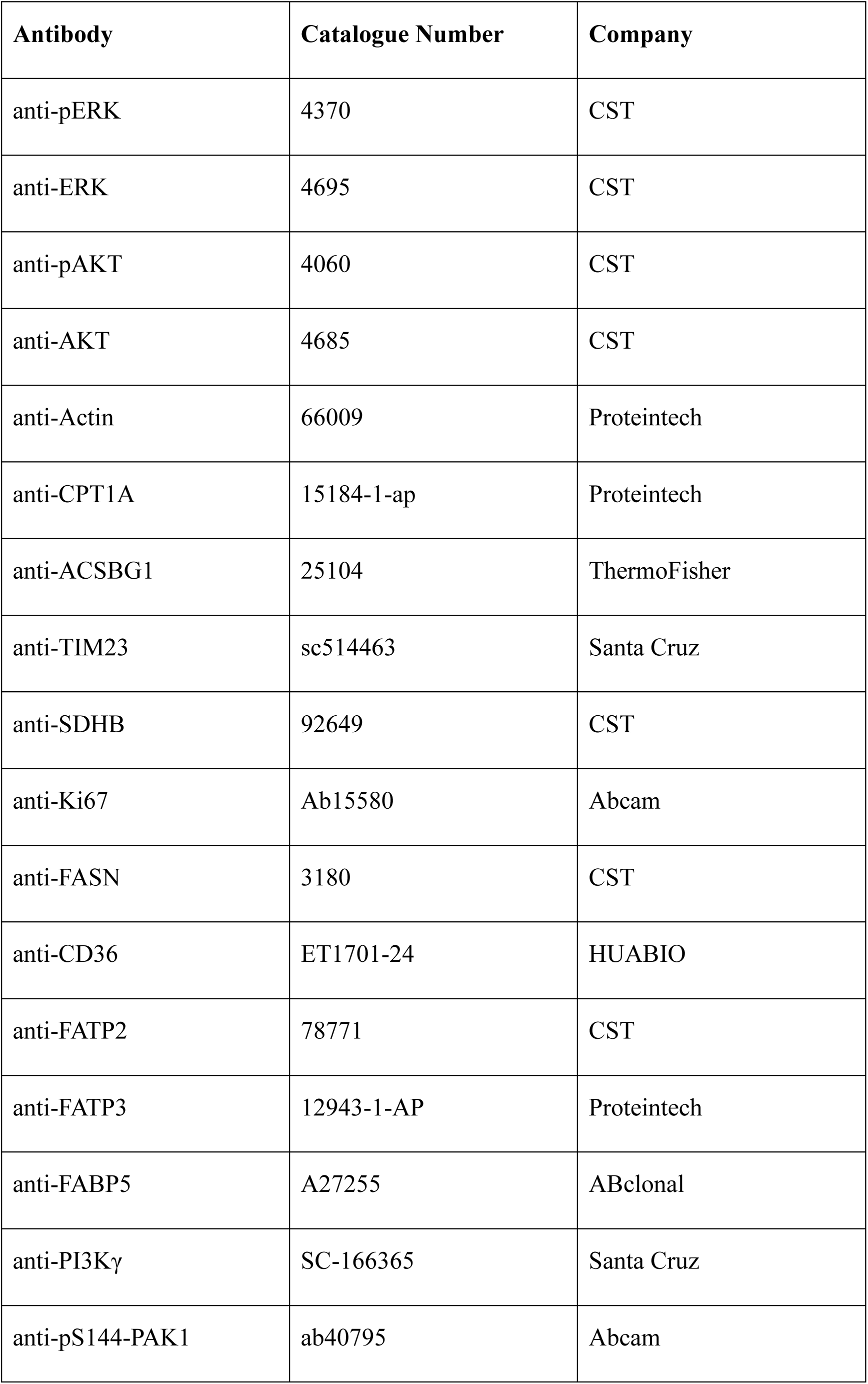
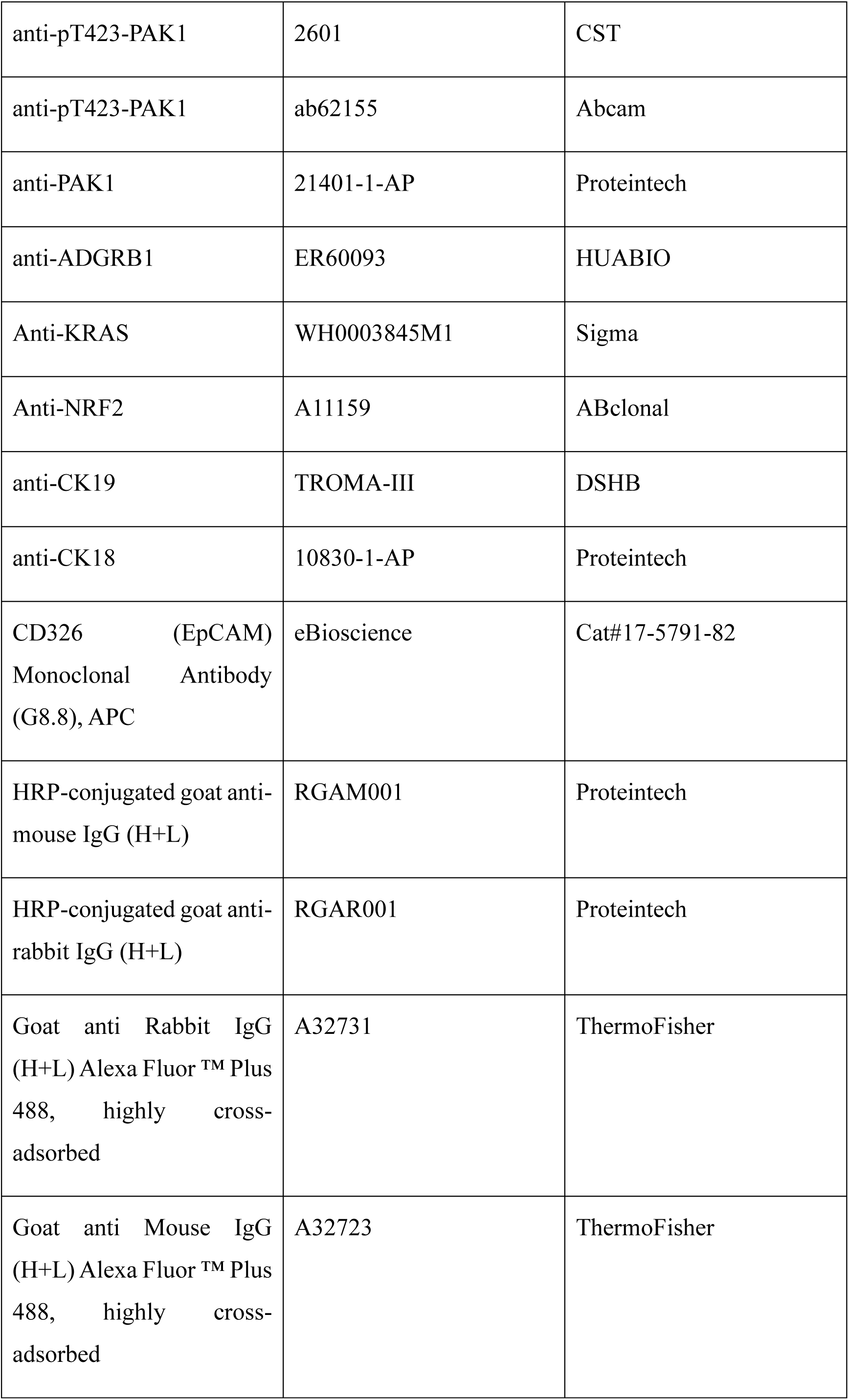
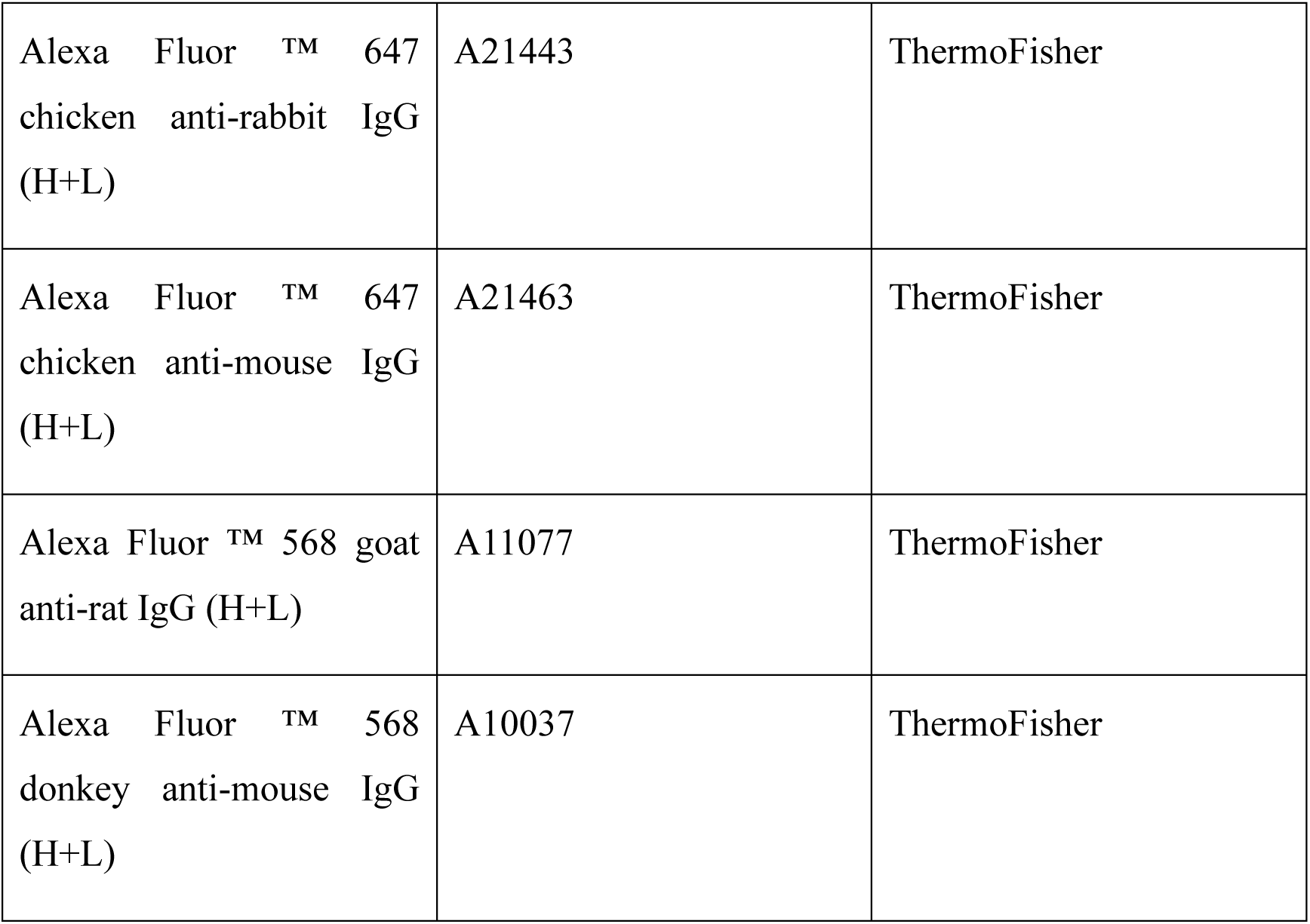
Antibody List.

**Table 2:**
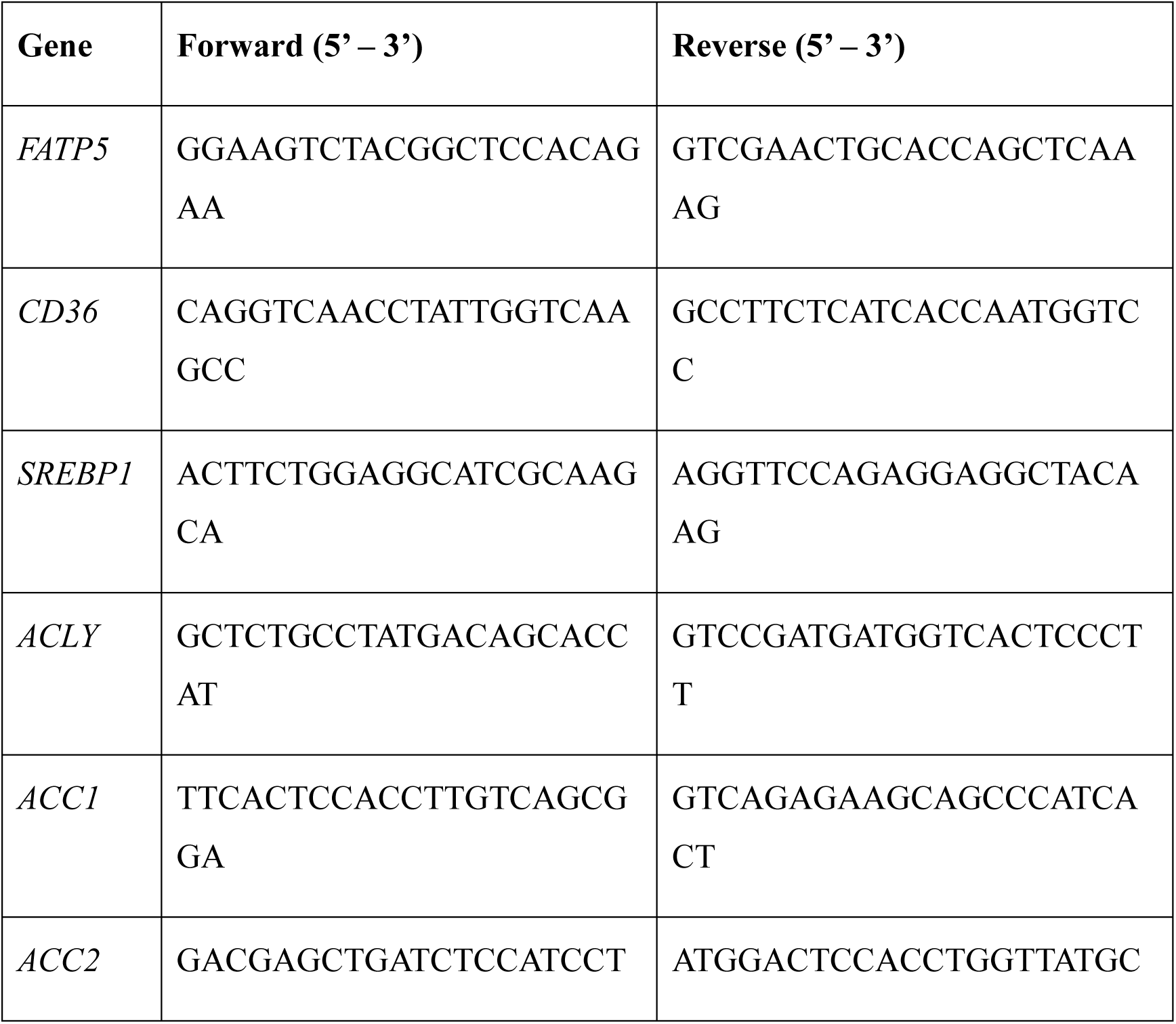
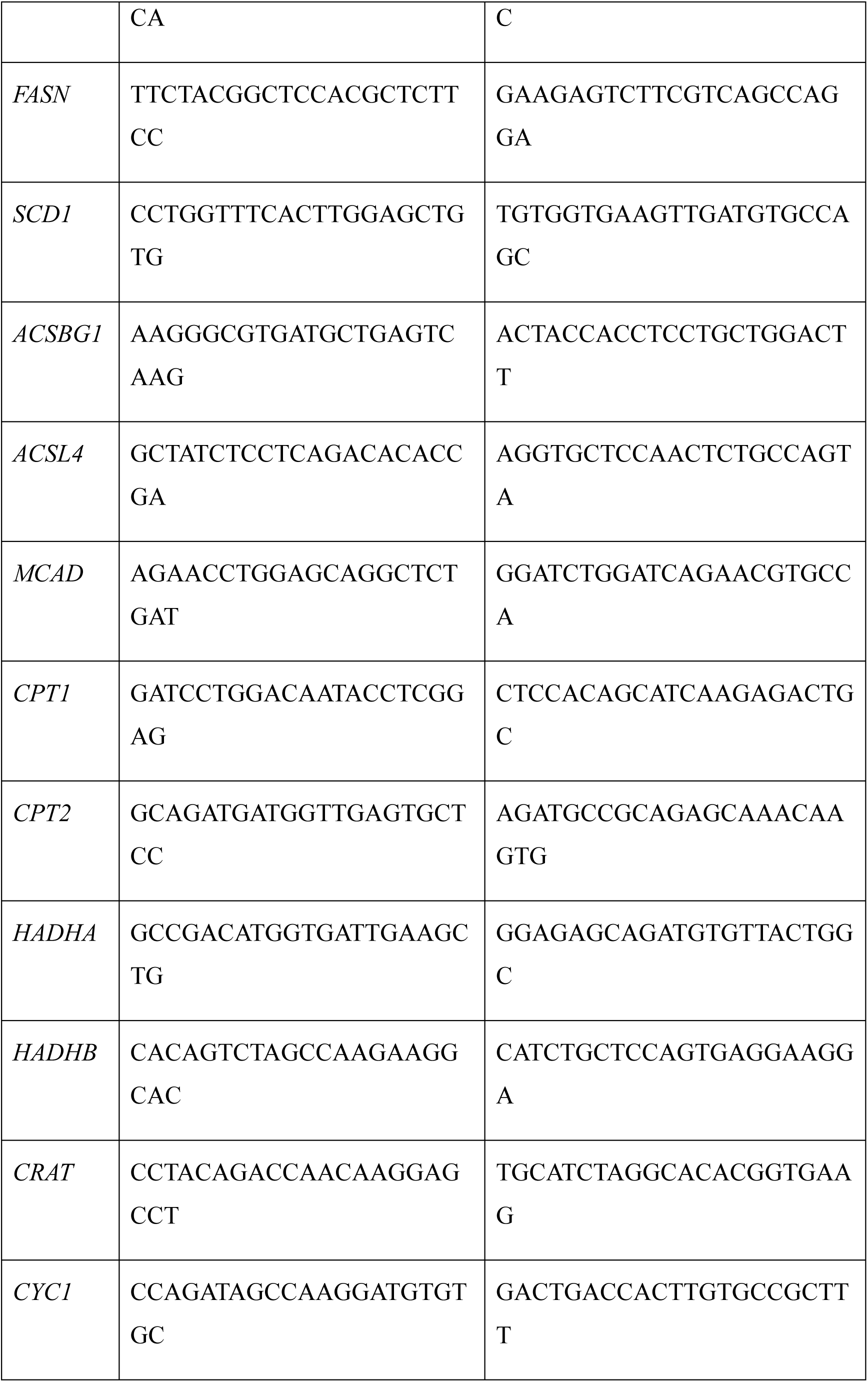
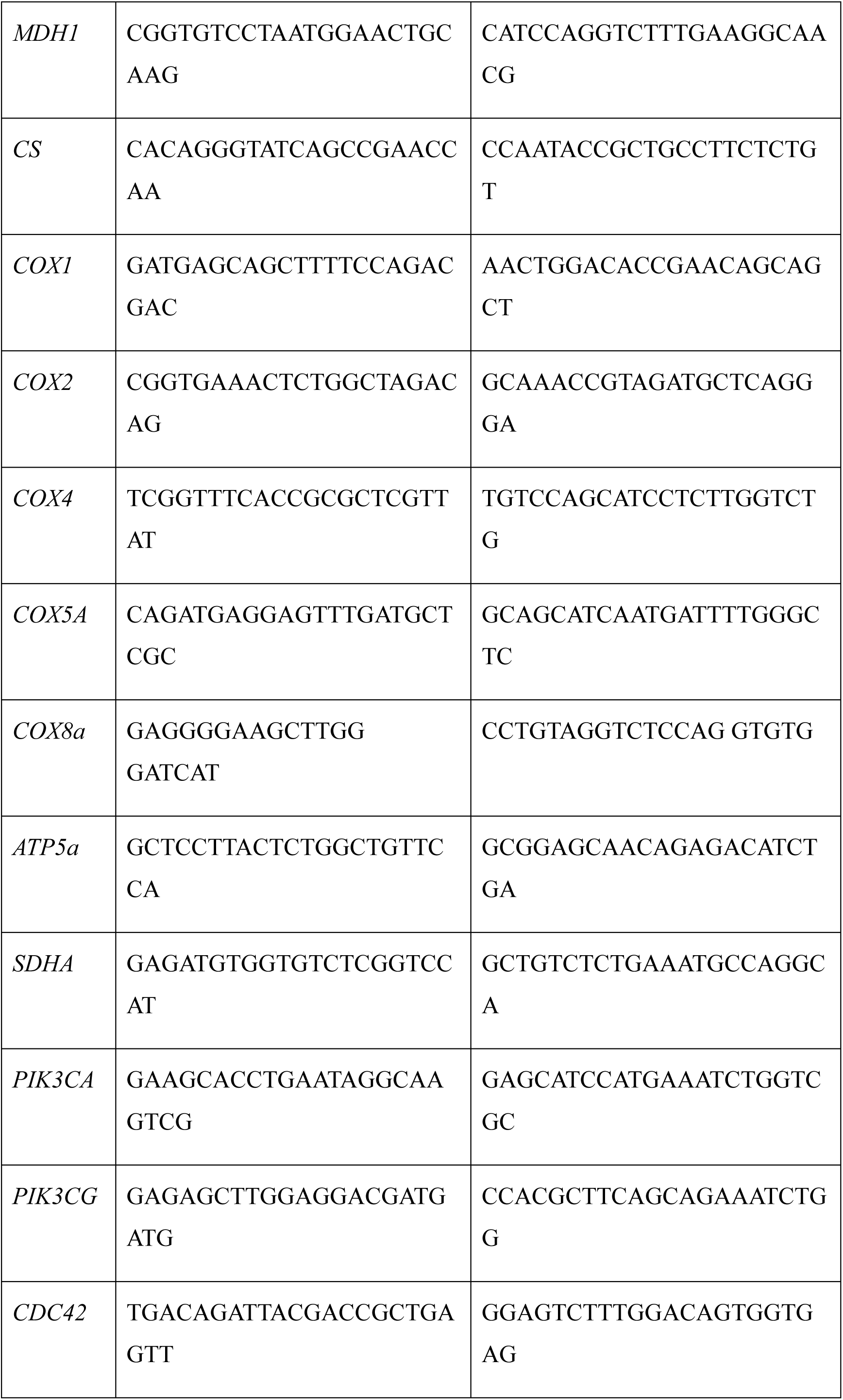
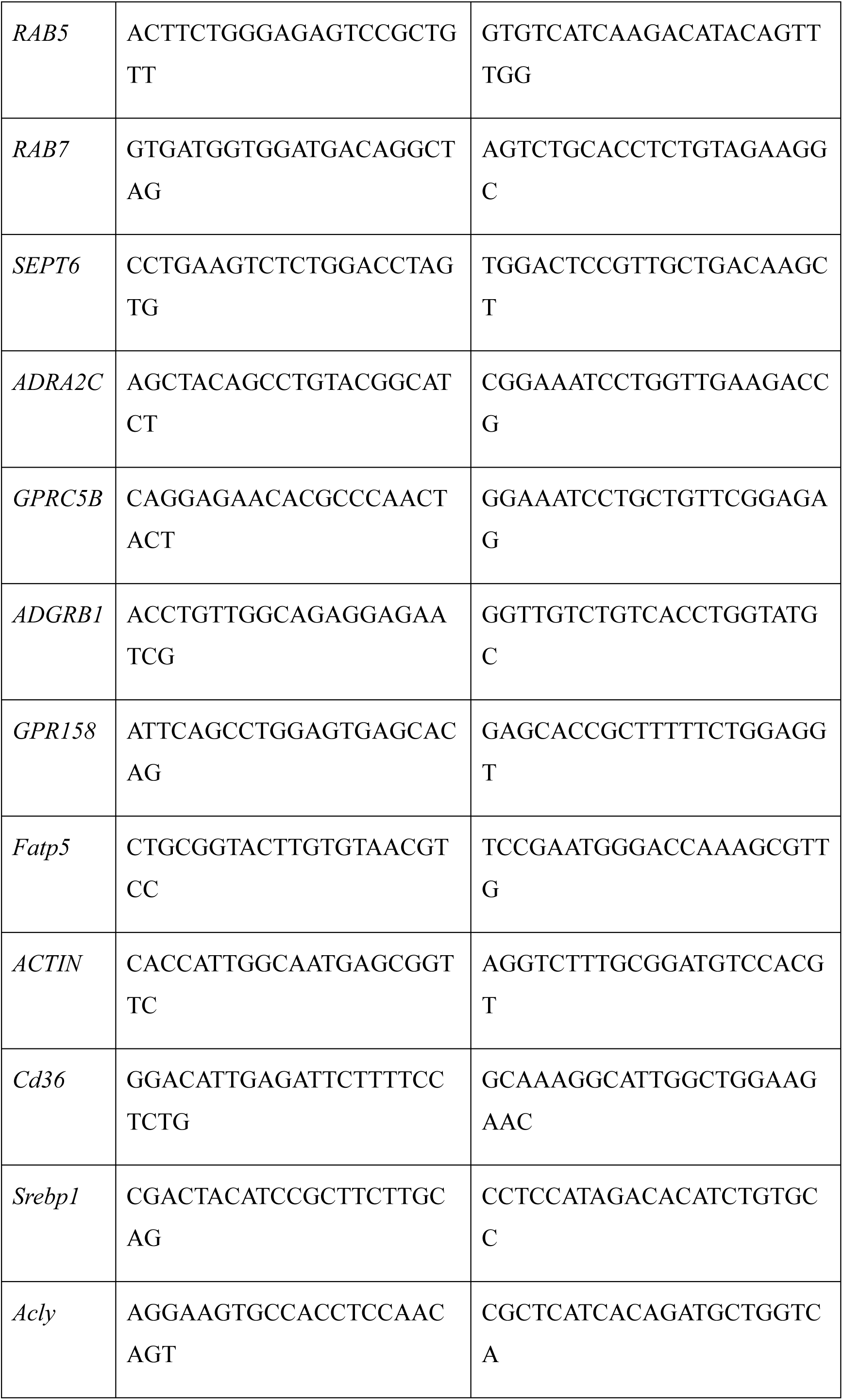
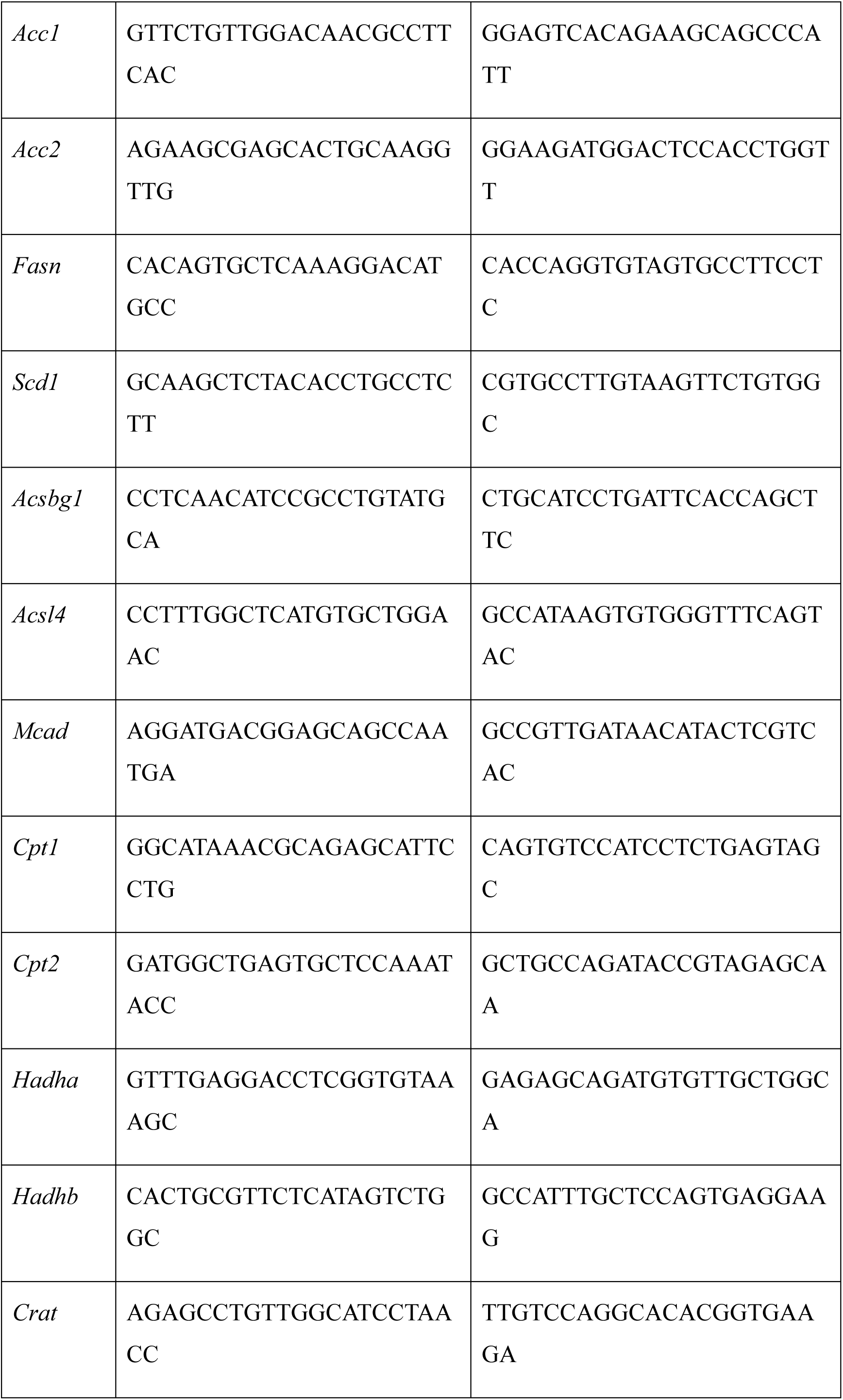
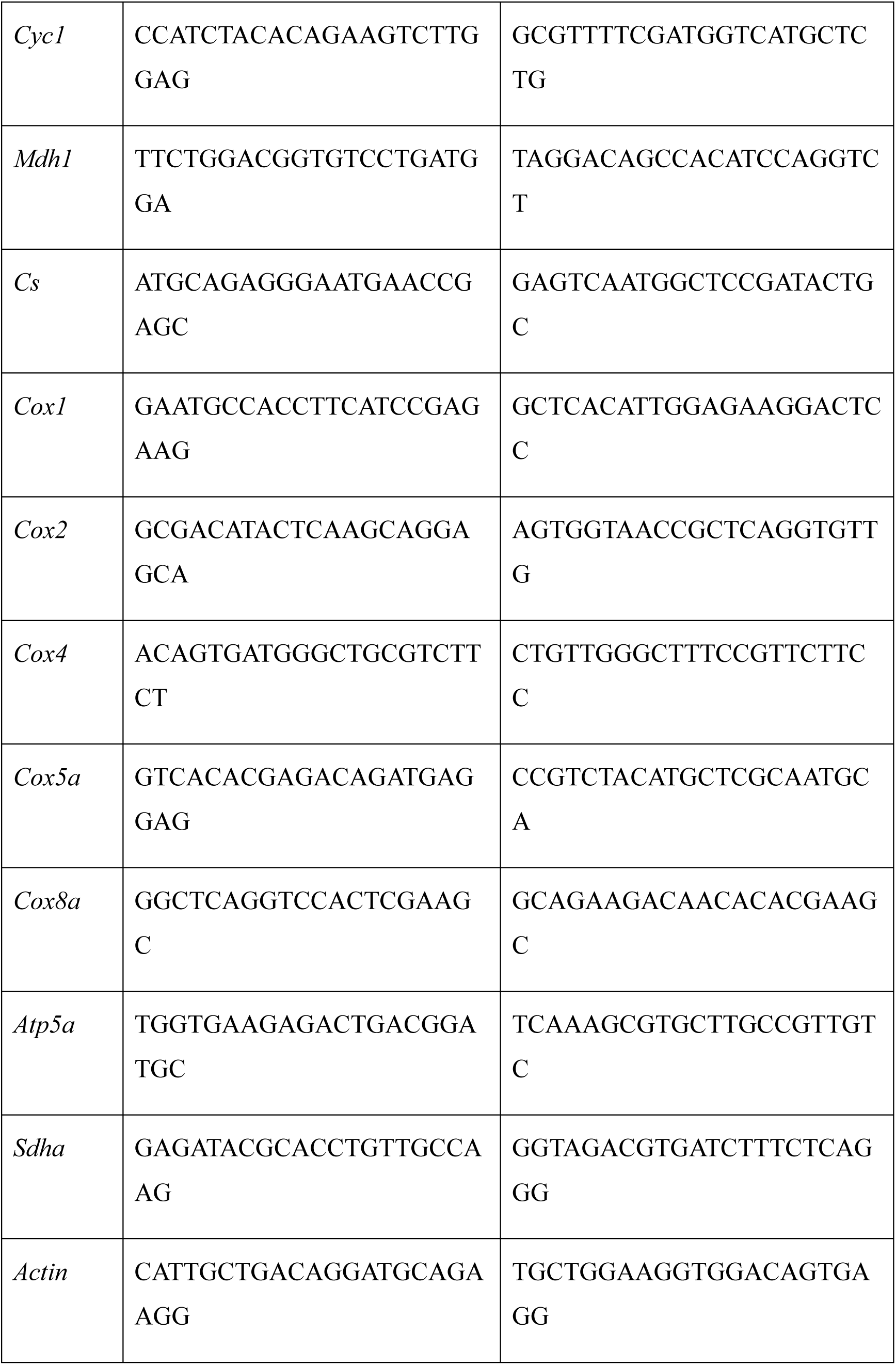
Real-time PCR Primer List.

## Acknowledgements

We thank members of the Dr. H.S. laboratory, Drs. Michael Karin and Andrew M Lowy for discussions; the Core Facility of Shanghai Medical College, Fudan University for microscopy services. Research was supported by grants from the National Natural Science Foundation of China (Grant Nos. 82372884, 82573758, 2022hwyq29 to H.S., 82372644 to F.Y., 82120108012 and 81930086 to B.S.); Noncommunicable Chronic Diseases-National Science and Technology Major Project (2026ZD0555800 to H.S.); the Shanghai Natural Science Foundation (Grant No. 23ZR1413600 to H.S.); the Fund of Fudan University and Cao’ejiang Basic Research (Grant No. 24FCA02 to H.S.); the Fund of Anhui Province Higher Education Outstanding Young Researcher (Grant No. 2024AH020007 to F.Y.); Clinical Research Special Project of Anhui Provincial Department of Science and Technology (202204295107020008 to B.S.); Research Program of Anhui Provincial Department of Education (B.S.); and the USA NIH (U01 CA274295-04 and R01 CA285997 to M.K.). Author contributions H.S. conceived the project. H.S., F.Y., and Z.Y. designed the study and Z.Y. and F.Y performed most experiments. F.Y. and Z.Y. performed IHC analysis of human and mouse PDAC. Z.Y., B.L., C.W., D.Z., and Z.M. performed immunoblotting and qPCR analysis. Z.Y., F.Y., C.W., Y.M., D.Z. and Z.M. performed orthotopic PDAC cell implantations. Z.Y., F.Y., C.W., and Z.M. prepared human PDAC organoids. Z.Y., G.W. performed the mass spectrometry imaging experiments. B.S. collected human PDAC tissue and supervised and supported F.Y. and Z.Y.. H.S., F.Y., and Z.Y. wrote the manuscript, with all authors contributing and providing feedback and advice.

---

**Extended Data Figure 1.**
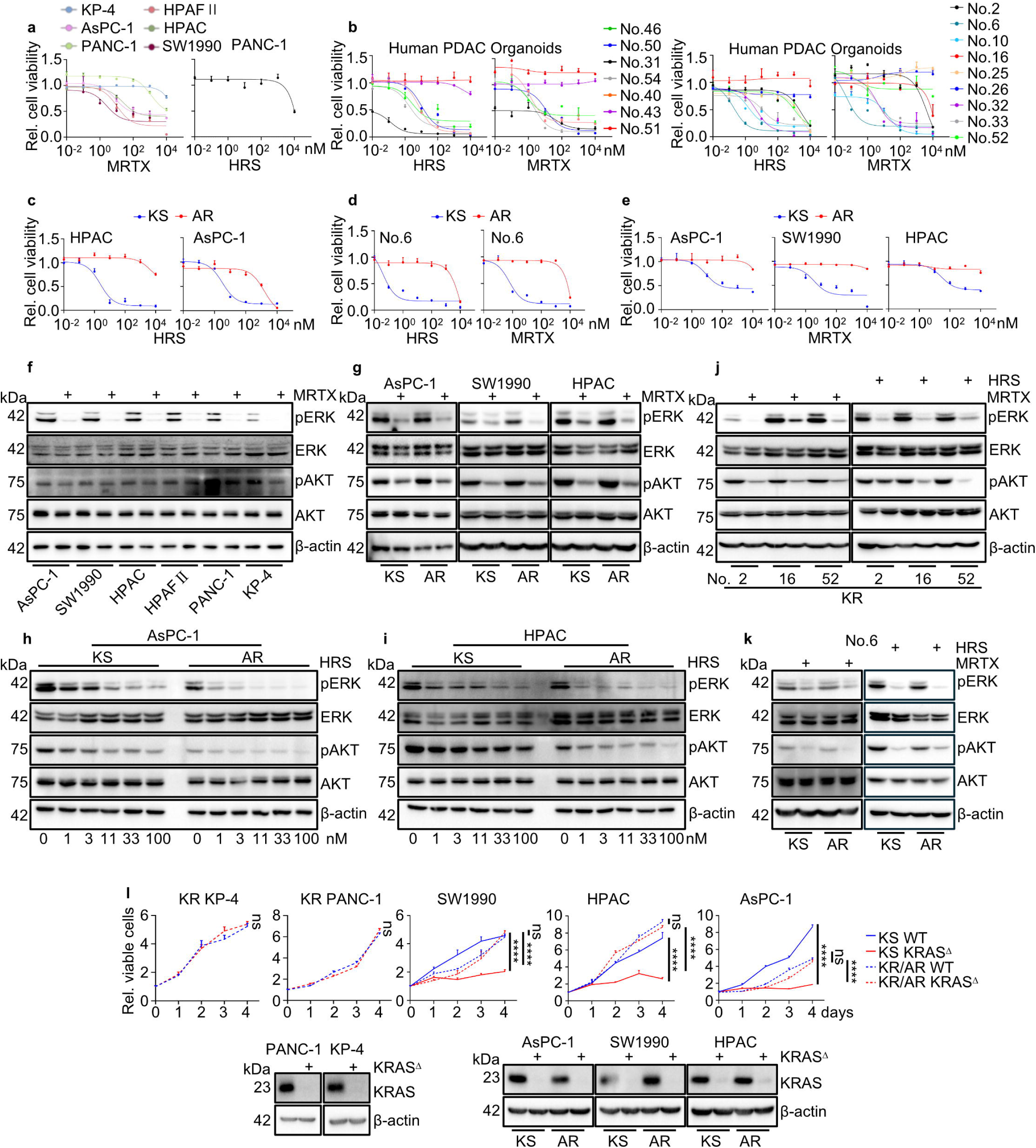
Profiling sensitivity and resistance to KRAS^G12D^i in human PDAC cells and organoids. **a-e,** The indicated *KRAS*^G12D^-mutant cells (**a**, **c**, **e**) and organoids (**b**, **d**) were treated with increasing concentration of MRTX1133 (MRTX) or HRS4642 (HRS) for 72 h (cells) or 7 days (organoids). Total viable cells are presented relative to vehicle (Veh). IC_50_ values are provided in Source Data. Parental HPAC, AsPC-1, SW1990, and No.6 cells are denoted as KS HPAC, AsPC-1, SW1990, and No.6, respectively. Acquired KRASi resistant HPAC, AsPC-1, SW1990, No.6 cells are labeled as AR. **f**-**i**, IB analysis of indicated proteins in human PDAC cells treated -/+ 100 nM MRTX for 2 h (**f**, **g**), or -/+ HRS at the indicated concentrations for 2 h (**h**, **i**). **j**, **k**, IB analysis of indicated proteins in human PDAC organoids treated -/+ 100 nM MRTX or HRS for 24 h. **l**, Viability of WT and KRAS-ablated (KRAS^Δ^) KS and KR/AR cells is presented relative to day 0. IB analysis of KRAS in the same cells is shown below. Data in (**a**-**e** and **l)** (n=3 independent experiments) are mean ± s.e.m. Statistical significance was determined using one-way ANOVA with Tukey post-hoc tests (SW1990, HPAC, and AsPC-1 in **l**) or two-sided unpaired t-test (KP-4, PANC-1 in **l**) based on data normality distribution. \**P* < 0.05, \*\**P* < 0.01, \*\*\**P* < 0.001, \*\*\*\**P* < 0.0001. NS, not significant. Exact *P* values are shown in Source Data.

**Extended Data Figure 2.**
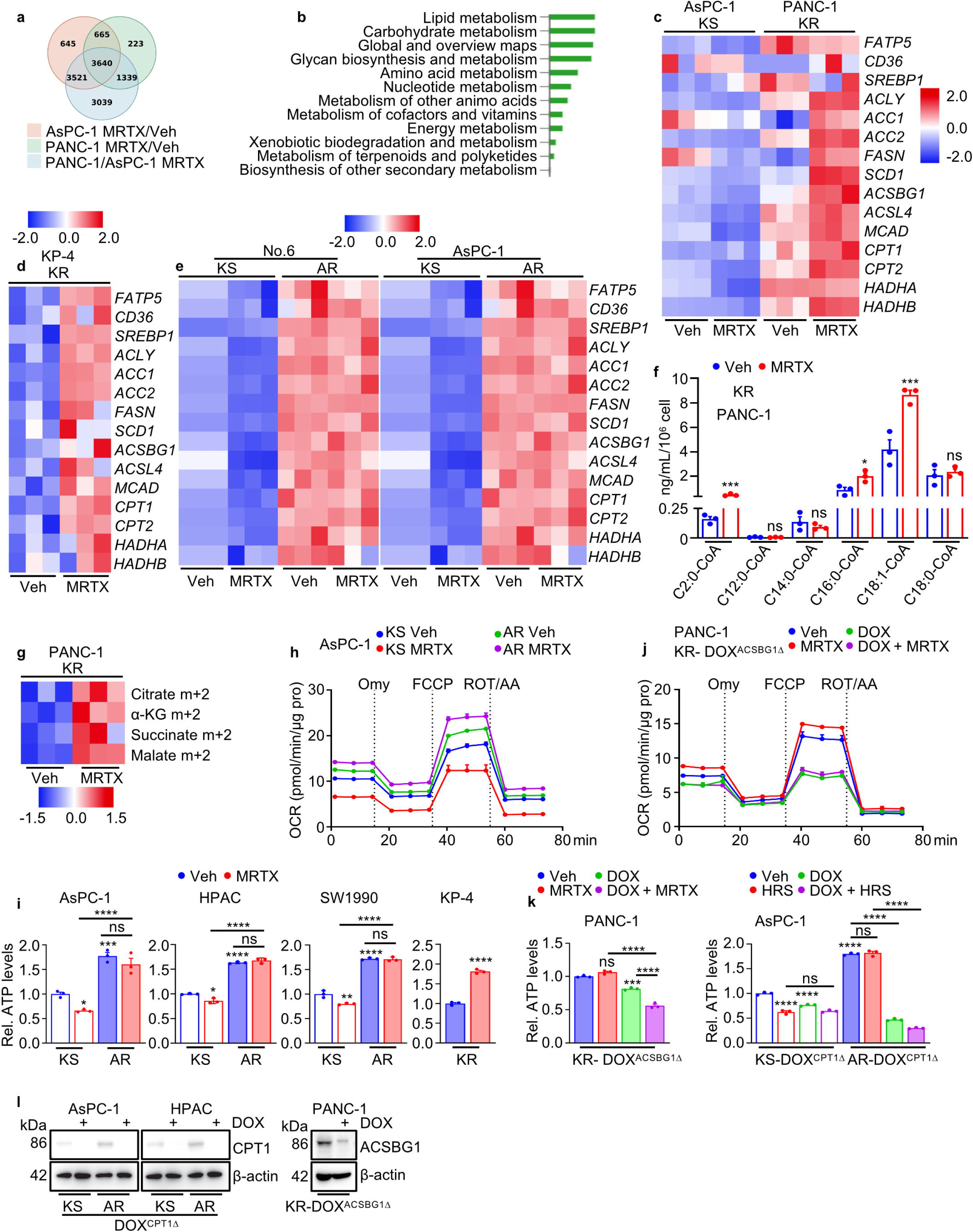
FA-driven oxidative phosphorylation is upregulated in KRASi-resistant cells. **a**-**c**, RNA-seq analysis was performed on KS (AsPC1) and KR (PANC-1) PDAC cells treated -/+ MRTX for 24 h. (**a**) Venn diagram showing the overlap of differentially expressed genes between MRTX- and Veh-treated cells and between MRTX-treated AsPC1 and MRTX-treated PANC-1 cells. (**b**) KEGG enrichment analysis of selected overlapping genes (n = 1,339 genes) from (**a**), representing genes whose expression is specifically altered in intrinsic resistant cells upon MRTX treatment. (**c**) Heatmap showing mRNA expression of lipid metabolism-related genes. Blue, replicates with low expression; red, replicates with high expression. **d**, **e**, qPCR analysis of indicated mRNAs in the indicated cell lines or organoids treated -/+ MRTX for 24 h. **f**, LC-MS quantification of intracellular acyl-coenzyme A levels in PANC-1 cells treated -/+ MRTX for 24 h. **g**, Fractional labelling of TCA cycle intermediates in PANC-1 cells cultured with [U-^13^C]-PA, -/+ MRTX for 12 h. **h**, OCR of KS and AR AsPC-1 cells treated -/+ MRTX for 24 h before and after treatment with Omy, FCCP, and rotenone/antimycin A (ROT/AA). **i**, Total cellular ATP in indicated KS and KR/AR cells treated -/+ MRTX for 48 h. Data are presented relative to untreated cells. KP-4 is intrinsically KRASi resistant (KR). **j**, OCR of KR PANC-1 cells expressing DOX^ACSBG1Δ^ treated -/+ DOX for 48h, followed with -/+ MRTX for 24 h before and after treatment with Omy, FCCP, and rotenone/antimycin A. **k**, Total cellular ATP in the indicated cells treated -/+ DOX for 48 h, followed with -/+ MRTX or HRS for 48 h. Data are presented relative to untreated cells. **l**, IB analysis of the indicated proteins in the indicated cells treated -/+ DOX for 48 h. Data in (**f**, **h**-**k**) (n=3 independent experiments) are mean ± s.e.m. Statistical significance was determined using two-sided unpaired *t*-test (**f**, **i** KP-4) or one-way ANOVA with Tukey post-hoc tests (**i**, **k**) based on data normality distribution. \**P* < 0.05, \*\**P* < 0.01, \*\*\**P* < 0.001, \*\*\*\**P* < 0.0001; ns, not significant. Exact *P* values are shown in Source Data.

**Extended Data Figure 3.**
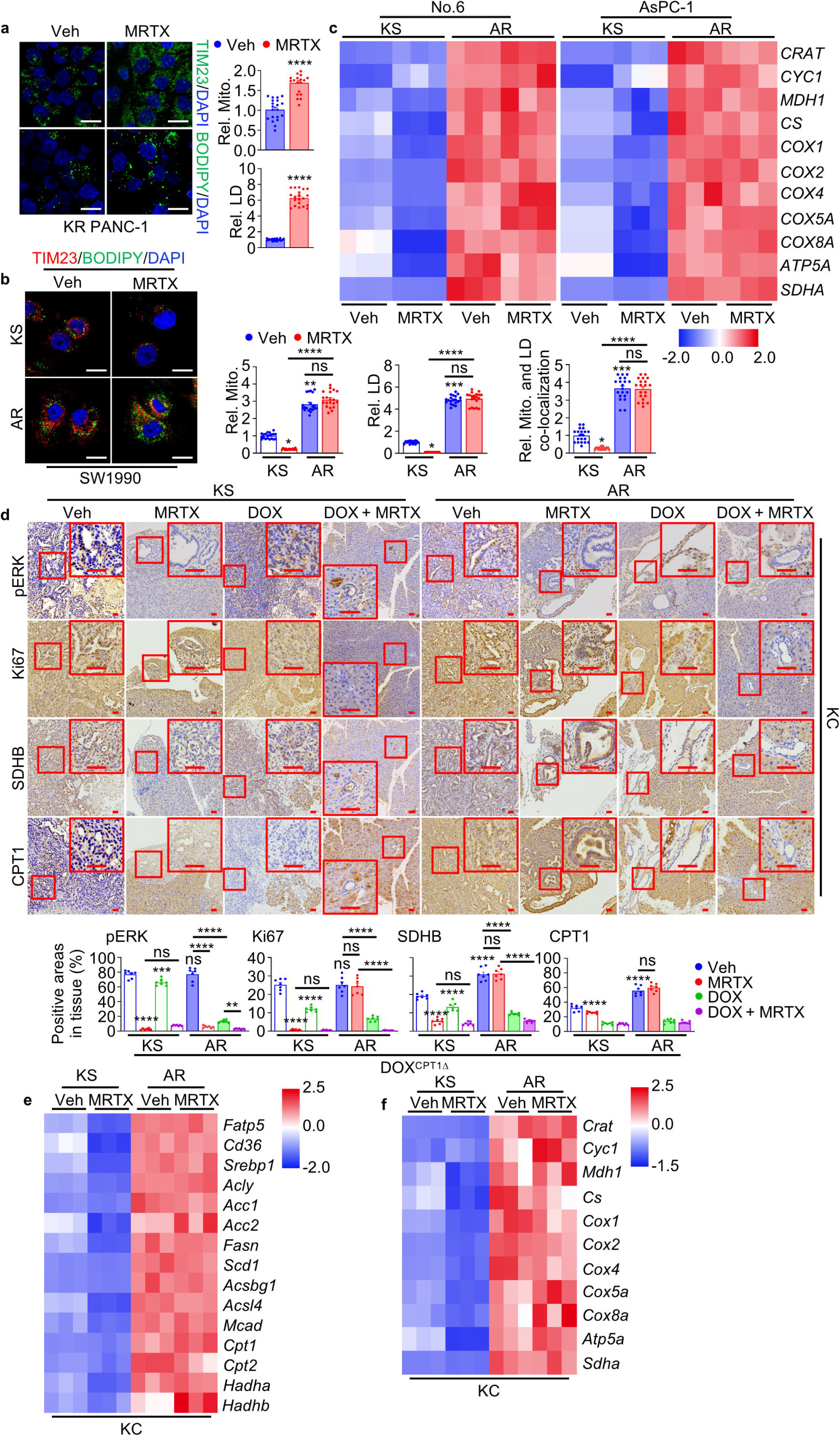
Mitochondrial FAO promotes KRASi resistance. **a**, **b**, Representative images, colocalization and quantification of mitochondria (TIM23) and LD (BODIPY) in PANC-1 cells (**a**), and KS and AR SW1990 cells (**b**), treated -/+ MRTX for 24 h. **c**, qPCR analysis of mitochondria-related mRNAs in KS and AR No.6 organoids and AsPC-1 cells treated -/+ MRTX for 24 h. **d**, IHC of pancreatic sections prepared 3 weeks after orthotopic transplantation of parental (KS) and acquired KRASi-resistance (AR) KC6141 cells expressing DOX^CPT1Δ^ into mice treated -/+ 30 mg/kg MRTX (i.p.), 0.2 mg/mL DOX (drinking water), or DOX + MRTX initiated 72 h post-transplantation. Boxed areas are further magnified. Image J determined staining intensity of indicated proteins in tissues is shown at the bottom. **e**, **f**, qPCR analysis of lipid metabolism-related (**e**) and mitochondria-related (**f**) mRNAs in the indicated tumors from (**d**). Data in (**a**, **b**) (n=20 fields) and (**d**) (n=7 fields) are mean ± s.e.m. Statistical significance was determined using two-sided unpaired t-test (**a**), Kruskal-Wallis test with Dunn post-hoc tests (**b**) or one-way ANOVA with Tukey post-hoc tests (**d**) based on data normality distribution. \**P*<0.05, \*\**P*<0.01, \*\*\**P*<0.001, \*\*\*\**P* < 0.0001. Exact *P* values are shown in Source Data. Scale bars (**a**, **b**) 20 μm, (**d**) 100 μm.

**Extended Data Figure 4.**
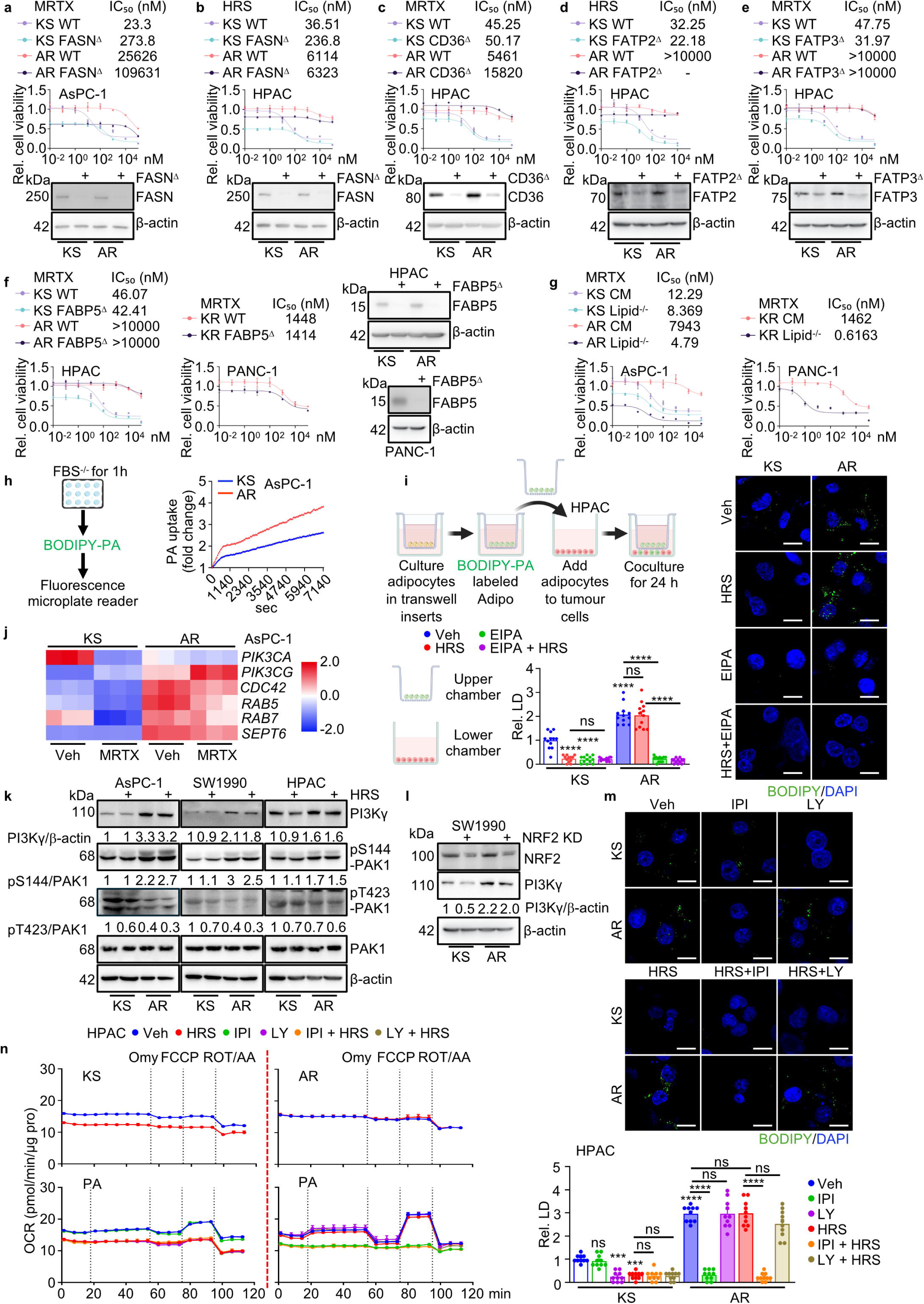
PI3Kγ is required for FAO in KRASi resistant cells. **a-f**, KS and AR AsPC-1, HPAC, or KR PANC-1 cells -/+ ablations of FASN, CD36, FATP2, FATP3, or FABP5 treated -/+ increasing concentrations of HRS or MRTX for 72 h. Cell viability is presented relative to untreated parental KS or KR (PANC-1) cells. IC_50_ values are presented on the top. IB analysis of FASN, CD36, FATP2, FATP3, and FABP5 expression is shown below or on the right. **g**, KS/AR AsPC-1 and KR PANC-1 cells cultured with completed (CM) or lipid-depleted media treated -/+ increasing concentrations of MRTX for 72 h. Cell viability is presented relative to CM- and vehicle-treated KS or KR (PANC-1) cells. IC_50_ values are presented on the top. **h**, Real-time lipid uptake assays were performed in KS and AR AsPC-1 cells after 1 h of serum starvation, followed by addition of BODIPY-PA for the indicated durations. BODIPY-PA uptake was quantified using a fluorescence microplate reader and is presented relative to the start of the assay. The experimental scheme is on the left. **i**, Adipocytes (3T3-L1) were preloaded with BODIPY-PA for 4 h in the upper chamber of a transwell plate, followed by coculture with KS and AR HPAC cells in the lower chamber, treated -/+ HRS, 25 μM EIPA, HRS + EIPA for 24 h. Representative images and quantification of BODIPY-LD are shown. The experimental scheme is on the top left. **j**, Heatmap showing MP-related mRNAs in KS and AR AsPC-1 cells treated -/+ MRTX for 24 h. Blue, replicates with low expression (z-score = −2); red, replicates with high expression (z-score = 2). **k**, KS and AR cells were treated -/+ HRS for 2 h followed by IB analysis of indicated proteins. **l**, IB analysis of indicated proteins in KS and AR SW1990 cells -/+ NRF2 KD. **m**, Representative images and quantification of BODIPY-LD in KS and AR HPAC cells cocultured with BODIPY-PA-preloaded 3T3-L1 adipocytes as in (**i**) and treated -/+ HRS, 1 μM LY294002 (LY), 1 μM IPI549 (IPI), HRS + IPI, or HRS + LY for 24 h. **n**, OCR of KS and AR HPAC cells incubated -/+ HRS, IPI, LY, HRS + IPI, or HRS + LY for 24 h before and after incubation -/+ PA, and treatment with Omy, FCCP, and rotenone/antimycin A. Data in (**a**-**g**, **h**, **n**) (n=3 independent experiments) and (**i**, **m**) (n=10-12 fields) are mean ± s.e.m. Statistical significance was determined using one-way ANOVA with Tukey post-hoc tests (**i**, **m**) based on data normality distribution. \**P* < 0.05, \*\**P* < 0.01, \*\*\**P* < 0.001, \*\*\*\**P* < 0.0001. Exact *P* values are shown in Source Data. Scale bars (**i**, **m**) 20 μm.

**Extended Data Figure 5.**
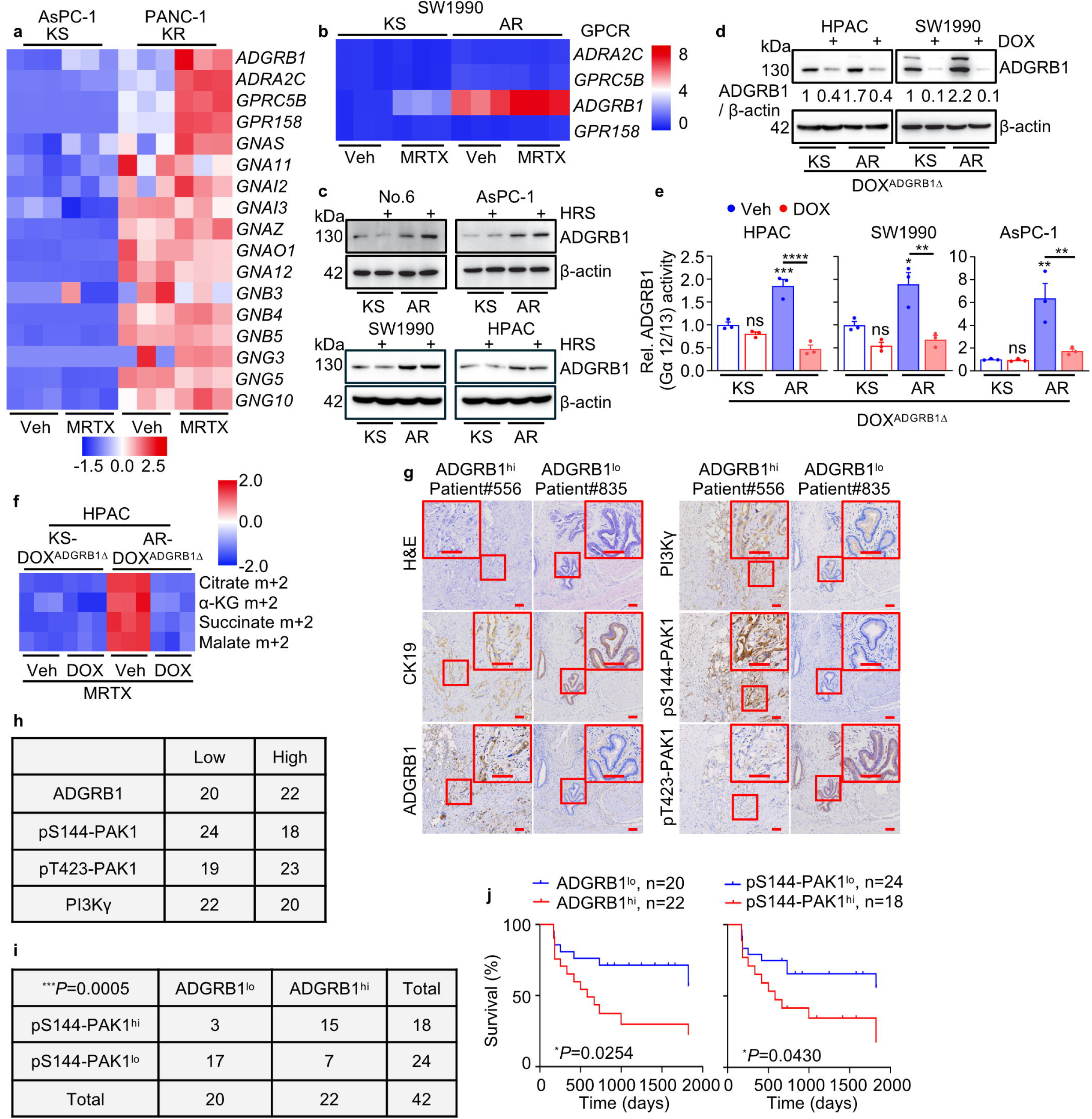
ADGRB1 sustains mitochondrial metabolism in KRASi-resistant tumor cells and predicts poor clinical outcome. **a**, **b**, Genes differentially expressed between KS (AsPC-1, SW1990) and KR/AR (PANC-1, SW1990) cells treated -/+ MRTX for 24 h. Blue, replicates with low expression; red, replicates with high expression. **c**, **d**, IB analysis of ADGRB1 in indicated organoids or cell lines treated -/+ HRS for 24 h (**c**) or -/+ DOX for 48 h (**d**). **e**, KS and AR cells expressing DOX^ADGRB1Δ^, SRF-responsive luciferase reporter (SRF-RE), and a Renilla-luciferase control were treated -/+ DOX for 48 h, followed by incubation with D-Luciferin. Normalized Firefly/Renilla luciferase ratios, reflecting ADGRB1 (Gα12/13) activity, are shown relative to untreated KS cells. **f**, Fractional labelling of TCA cycle intermediates in KS and AR HPAC cells expressing DOX^ADGRB1Δ^ treated -/+ DOX for 48 h, and then incubated with [U-^13^C]-PA and MRTX for 12 h. Blue, replicates with low expression; red, replicates with high expression. **g**, Representative IHC of resected ADGRB1^hi^ (#556) and ADGRB1^lo^ (# 835) human PDAC tissues. Boxed areas are further magnified. **h**, Numbers of human PDAC specimens (n = 42) positive for the indicated proteins, indicated as low and high expression. **i**, Correlation between expression levels of the indicated proteins in human specimens examined by a two-tailed Chi-square test. **j**, Comparisons of overall survival between patients with PDAC stratified according to ADGRB1 and pS144-PAK1. Significance was determined by log-rank test. Data in (**e**) (n=3 independent experiments) are mean ± s.e.m. Statistical significance was determined using one-way ANOVA with Tukey post-hoc tests (**e**). \**P* < 0.05, \*\**P* < 0.01, \*\*\**P* < 0.001, \*\*\*\**P* < 0.0001. Exact *P* values are shown in Source Data.

**Extended Data Figure 6.**
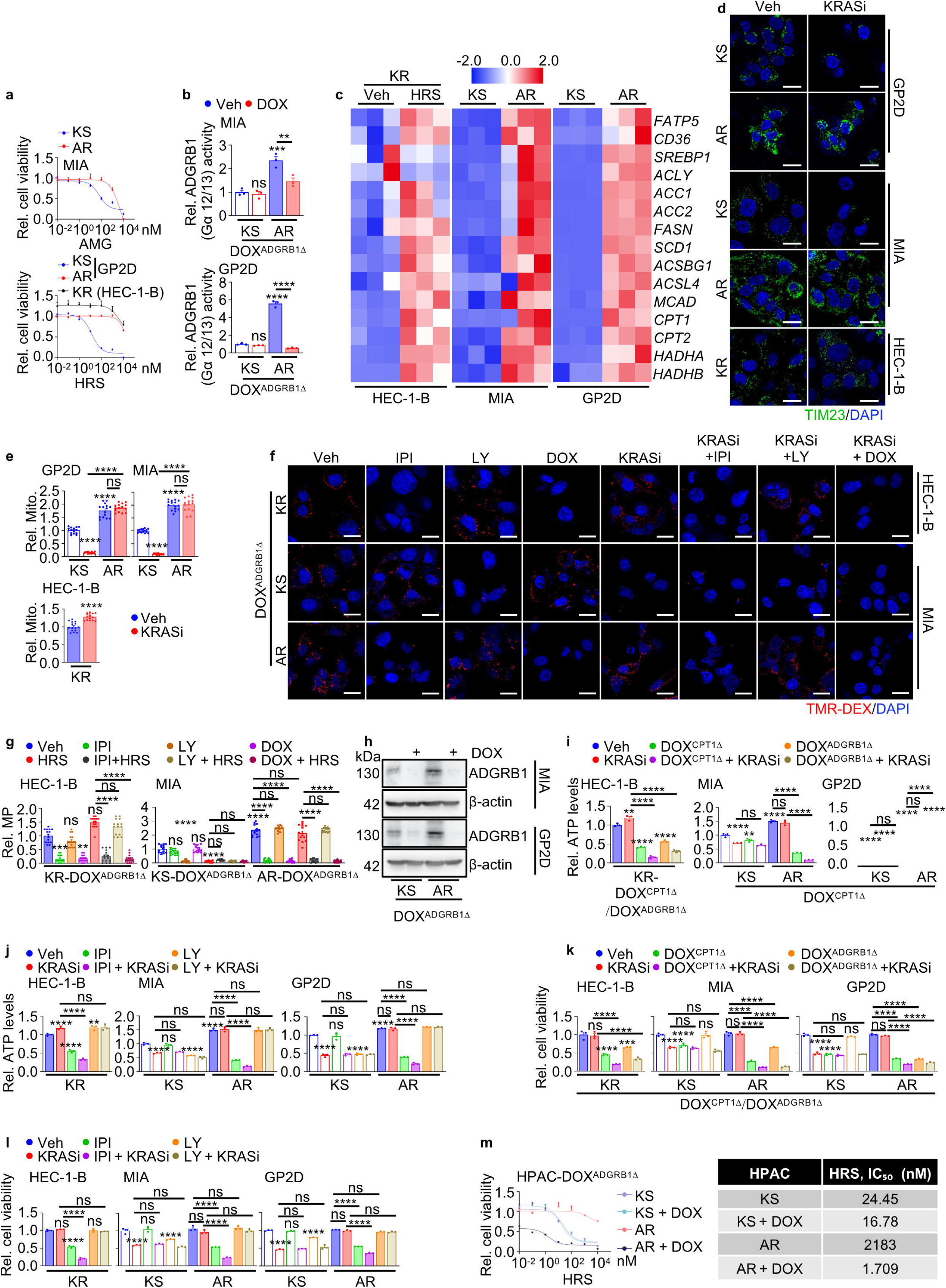
ADGRB1 signaling maintains MP, mitochondria and ATP content in KRASi-resistant cells across diverse KRAS-mutant cancers. **a**, The indicated cells were treated -/+ increasing concentrations of AMG510 (AMG) or HRS for 72 h. Cell viability is presented relative to untreated KS or KR (HEC-1-B) cells. IC_50_ values are presented in the Source Data. MIA, MIA-PaCa-2. **b**, The indicated KS and AR cells expressing DOX^ADGRB1Δ^, SRF-RE, and a Renilla-luciferase control were treated -/+ DOX for 48 h, followed by incubation with D-Luciferin. Normalized Firefly/Renilla luciferase ratios, reflecting ADGRB1 (Gα12/13) activity, are shown relative to untreated KS cells. **c**, Heatmap showing FA metabolism-related mRNAs in the indicated cells treated -/+ HRS for 24 h. Blue, replicates with low expression; red, replicates with high expression. **d**, Representative images of mitochondria (TIM23) in indicated cells treated -/+ KRASi (AMG for MIA, or HRS for GP2D and HEC-1-B) for 24 h. **e**, Quantification of mitochondrial number in (**d)**. **f**, Representative images of MP in the indicated cells incubated -/+ DOX for 48h, followed by treatment -/+ IPI, LY, HRS, HRS + IPI, or HRS + LY for 24 h. **g**, Quantification of MP in (**f**). **h**, IB analysis of ADGRB1 in the indicated cells incubated -/+ DOX for 48 h. **i**-**l**, The indicated cells were treated -/+ the indicated chemicals for 48 h. Total cellular ATP (**i**, **j**) or cell viability (**k**, **l**) are presented relative to untreated KS or KR (HEC-1-B) cells. KRASi indicates AMG for MIA, or HRS for HEC-1-B and GP2D. **m**, KS and AR HPAC expressing DOX^ADGRB1Δ^ and treated -/+ DOX for 48 h, followed by treatment with increasing concentrations of HRS for 72 h. Cell viability is presented relative to untreated KS cells. IC_50_ values are shown to the right. Data in (**a**, **b**, **i**-**m**) (n=3 independent experiments) and (**e**, **g**) (n=15 fields) are mean ± s.e.m. Statistical significance was determined using one-way ANOVA with Tukey post-hoc tests (**b**, **e**, **g**, **i**-**l**), two-sided unpaired t-test (**e** HEC-1-B) and Kruskal-Wallis test with Dunn post-hoc tests (**g** HEC-1-B) based on data normality distribution. \**P* < 0.05, \*\**P* < 0.01, \*\*\**P* < 0.001, \*\*\*\**P* < 0.0001. Exact *P* values are shown in Source Data. Scale bars (**d**, **f**) 20 μm.

**Extended Data Figure 7.**
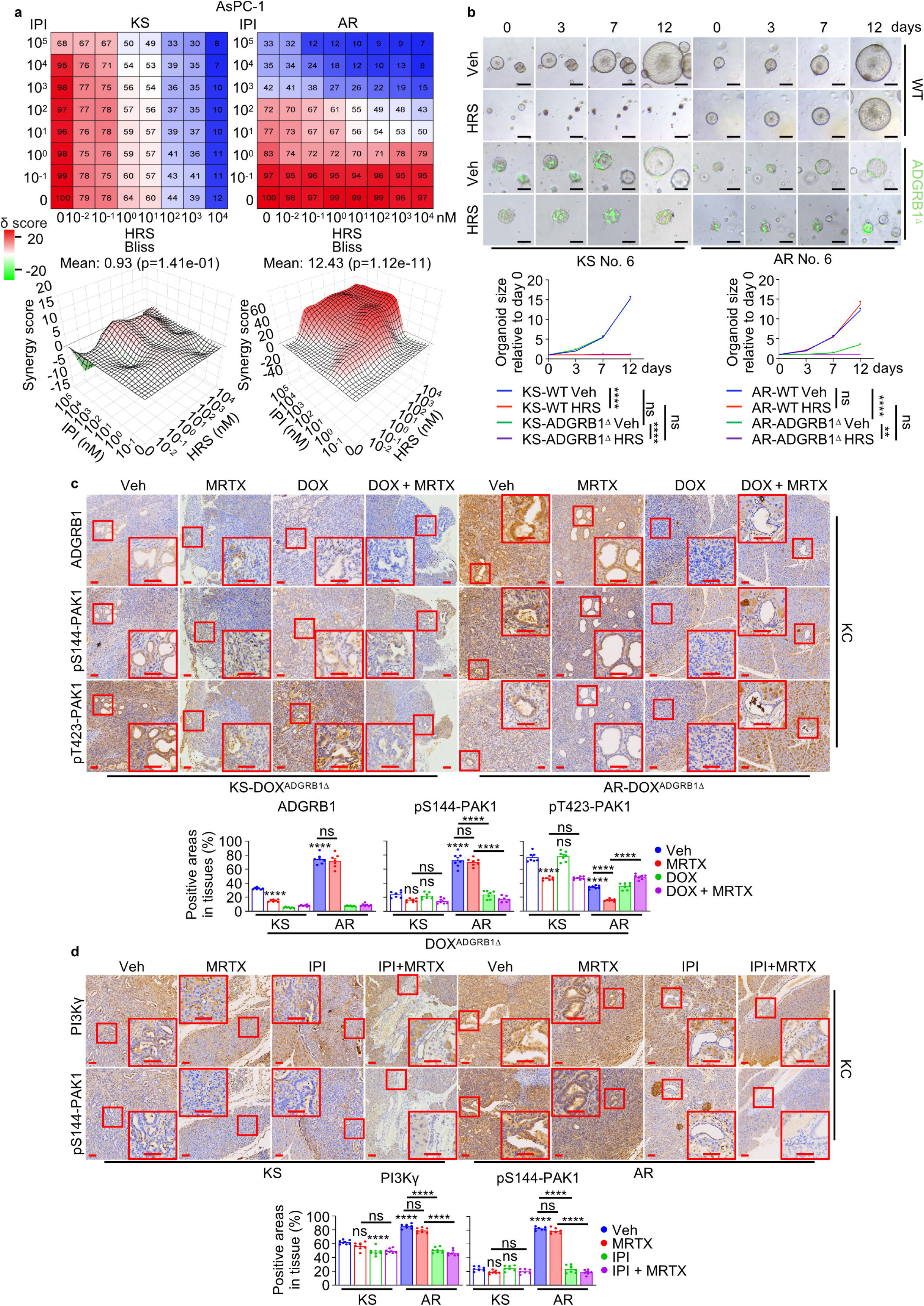
Inhibition of the ADGRB1-PI3Kγ axis selectively suppresses KRASi resistance. **a**, KS and AR AsPC-1 cells were treated with IPI and/or HRS at the indicated concentrations for 3 days and analyzed for cell viability by CCK8 assay. Upper is dose-response matrix (viability) for IPI and HRS. Blue intensity indicates the degree of inhibition. Lower are the 3D synergy plots generated by SynergyFinder^+^ software. The synergy score is shown by red (>0) and green (<0). The synergy score > 10 indicates strong synergic effect. **b**, Representative images and quantification of growth of KS and AR No.6 organoids expressing -/+ GFP-tagged ADGRB1-ablated vector (ADGRB1^Δ^), and treated -/+ HRS for 12 days. Individual size of organoid is presented relative to day 0. **c**, Representative IHC and quantification of indicated proteins in pancreata 3 weeks after orthotopic transplantation of indicated cells into mice followed by treatment -/+ MRTX, DOX, or DOX + MRTX, initiated 72 h post-transplantation. Boxed areas are further magnified. **d**, Representative IHC and quantification of indicated proteins in pancreata 3 weeks after orthotopic transplantation of the indicated cells into mice and treatment -/+ MRTX, 15 mg/kg IPI (p.o.), or MRTX + IPI, initiated 72 h post-transplantation. Boxed areas are further magnified. Data in (**b**) (n= 10 organoids) and (**c**, **d**) (n=7 fields) are mean ± s.e.m. Statistical significance was determined using one-way ANOVA with Tukey post-hoc tests (**b**-**d**) based on data normality distribution. \**P* < 0.05, \*\**P* < 0.01, \*\*\**P* < 0.001, \*\*\*\**P* < 0.0001. Exact *P* values are shown in Source Data. Scale bars (**b**) 100 μm, (**c**, **d**) 100 μm.

## Methods

### Cell culture

All cells were incubated at 37 °C in a humidified chamber with 5% CO2. AsPC-1, PANC-1, HPAC, SW1990, GP2D, MIA PaCa-2 (MIA), HEC-1-B, and UN-KC-6141 (KC) cells were maintained in Dulbecco’s modified Eagle’s medium (DMEM) (Gibco) supplemented with 10% fetal bovine serum (FBS) (Gibco). KP-4 was maintained in Roswell Park Memorial Institute 1640 medium (1640) (Gibco) supplemented with 10% FBS. HPAF-II was maintained in Minimum Essential Medium (MEM) (Gibco) supplemented with 10% FBS and 1% non-essential amino acids (NEAA) (Gibco). 3T3-L1 cells were maintained in DMEM supplemented with 10% Calf Serum (Procell). MIA was purchased from ATCC. PANC-1, HPAC, SW1990 and GP2D were purchased from Yuchun Biology (Shanghai, China). KP-4 was purchased from Meisen Biology (Shanghai, China). 3T3-L1 and HEC-1-B were purchased from Biowing Biology (Shanghai, China). HPAF-II was purchased from Zhongqiaoxinzhou Biotechnology (Shanghai, China). AsPC-1 and HEK293T were gifts from Dr. Michael Karin (Sanford Burnham Prebys). KC cells were provided by Dr. Surinder K. Batra (University of Nebraska Medical Center). All media were supplemented with penicillin (100 U/mL) and streptomycin (100 μg/mL). All cells were partially authenticated by visual morphology and by assessing the expression of downstream KRAS signaling proteins (pERK and pAKT) in response to allele-specific KRAS inhibitors. All cell lines were routinely tested for mycoplasma contamination.

### Generation of drug-resistant cell lines and organoids

To generate AR cells, KS AsPC-1, HPAC, SW1990, GP2D, and KC were seeded at 50% confluency and continuously treated with 100 nM MRTX1133 (MRTX) or HRS-4642 (HRS). Culture medium was refreshed every 2 days, and cells were passaged upon reaching full confluency. After at least three passages, the drug concentration was increased to 500 nM and subsequently maintained throughout culturing. For all assays, a 72-h drug washout period was applied prior to further analysis. AR cells were considered successfully established when the IC_50_ of MRTX or HRS increased by at least 50-fold compared with KS cells. For KR MIA-PaCa-2 cells, AMG510 was used in KS cells, following the same procedure.

To generate AR organoids, KS organoids were initially treated with 10 nM MRTX until organoids reached a diameter of approximately 300 μm. After passaging, organoids were cultured with 100 nM MRTX until reaching a similar size. This stepwise escalation was repeated until organoids were able to proliferate under 500 nM MRTX treatment. Resistance was defined as a ≥50-fold increase in IC_50_ relative to KS organoids. A 72-h drug washout period was applied before all assays.

### Plasmids

For gene ablations, Tet-pLKO.1-puro-mouse (m) Cpt1 (TRCN0000110598), Tet-pLKO.1-puro-human (h) CPT1 (TRCN0000036279), Tet-pLKO.1-puro-hACSBG1 (TRCN0000150785), pLKO.1-puro-hFASN (TRCN0000003127), pLKO.1-puro-hCD36 (TRCN0000056999), pLKO.1-puro-hFATP2 (TRCN0000043389), pLKO.1-puro-hFATP3 (TRCN0000043410), pLKO.1-puro-hFABP5 (TRCN0000011895), pLKO.1-puro-hKRAS (TRCN0000010369), pLKO.1-puro-hGPRC5B (TRCN000000 8982), pLKO.1-puro-hGPR158 (TRCN0000062923), pLKO.1-puro-hADRA2C (TRCN0000008075) and pLKO.1-puro NRF2 (TRCN0000007558) were obtained from Sigma. Tet-pLKO-puro-mAdgrb1 (GCACCCTTGGAGATCGAGTTTG) and Tet-pLKO-puro-hADGRB1 (GAGTTTGCCCACATGTATAAT) were obtained from VectorBuilder. pX330-ADGRB1-EGFP was obtained from Thermo Fisher (CRISPR1126533_SGM). pGL4.34 (luc2P SRF-RE Hygro) and pRL-TK Renilla were obtained from Promega. pLVX-IRES-BSD-PI3Kγ^R1021C^-Flag and pLV3-mKi67-Akaluciferase were obtained from Sangon Biotech.

### Stable cell line construction

Lentiviral particles were generated as before^39^. KS/AR/KR AsPC-1, PANC-1, HPAC, SW1990, GP2D, MIA PaCa-2, HEC-1-B, KP-4, and KC6141 cells were transduced by combining 1 mL of viral particle-containing medium with 8 μg/mL polybrene. The cells were fed 8 h later with fresh medium, and selection was initiated 48 h after transduction using 1.25 μg/mL puromycin or 10 μg/mL blasticidin. For the Tet-on system, doxycycline (DOX) was used at a concentration of 2 μg/mL.

### Mice

C57BL/6, *Adipoq*-CreERT2, and *Atgl*/*Pnpla2^f/f^*mice were purchased from Jiangsu GemPharmatech. *Adipoq*-CreERT2 and *Pnpla2^f/f^*mice were interbred to obtain the compound mutants *Adipoq*-CreERT2;*Pnpla2^f/f^* (termed *Atgl*^Δadipo^). For ablation of ATGL in adipocytes, 6 weeks old *Atgl*^Δadipo^ mice were treated with tamoxifen (75 mg/kg, dissolved in corn oil) via i.p. once daily for 5 consecutive days. Experiments were performed at least two weeks after the final tamoxifen injection. Mice matched for age and gender were randomly allocated to different experimental groups based on their genotypes. No sample size pre-estimation was performed, but as many mice per group as possible were used to minimize type I/II errors. Both male and female mice were used unless otherwise stated. Blinding of mice was not performed except for IHC analysis. All mice were maintained in filter-topped cages on autoclaved food and water at constant temperature and humidity in a pathogen-free controlled environment (23 ± 2°C, 50-60%) with a standard 12 h light/12 h dark cycle. Experiments were performed in accordance with Fudan University Institutional Animal Care and Use Committee guidelines and regulations. Dr. Hua Su’s Animal Protocol DSF-2023-003 was approved by the Fudan University Institutional Animal Care and Use Committee. The number of mice per experiment and are indicated in the figure legends.

### Orthotopic PDAC cell implantation

KS/AR KC, KC-DOX^CPT1Δ^, or KC-DOX^ADGRB1Δ^ cells were orthotopically injected into three-month-old C57BL/6, *Atgl*^WT^, and *Atgl*^Δadipo^ mice as described^39^. Following surgery, mice were given buprenorphine subcutaneously at a dose of 0.05-0.1 mg/kg every 4-6 h for 12 h and then every 6-8 h for 3 additional days. For ablation of CPT1 or ADGRB1, mice were provided with drinking water containing 200 μg/mL DOX (Sangon Biotech), starting 72 h post-cell transplantation. DOX-containing water was replaced with freshly prepared solution every 2 days. MRTX1133 (MCE) was administered via intraperitoneal injection (i.p.) at a dosage of 30 mg/kg daily in 10% (w/v) 2-hydroxypropyl-β-cyclodextrin (HP-β-CD) vehicle. IPI549 (Selleck) was administered by oral gavage (p.o.) at a dosage of 15 mg/kg daily in a vehicle consisting of 10% DMSO / 35% Tween-80 / 55% H_2_O (v/v/v). EIPA (Selleck) was administered by i.p. at 30 mg/kg every other day in 5% DMSO / 5% Tween-80 / 35% PEG-300 / 55% saline (v/v/v/v). HRS-4642 (Selleck) was administered by i.p. at 10 mg/kg daily in the same vehicle. All drug treatments were initiated 72 h after transplantation. Mice were analyzed 3 weeks after transplantation.

### Human specimens

42 human PDAC specimens bearing *KRAS*^G12D^ mutation were acquired from patients who were diagnosed with PDAC between January 2017 and May 2023 at The First Affiliated Hospital of Anhui Medical University (Hefei, Anhui, China). All patients received standard surgical resection without chemotherapy before surgery. Paraffin-embedded tissues were processed by a pathologist after surgical resection and confirmed as PDAC before further investigation. Overall survival duration was defined as the time from the date of diagnosis to that of death or last known follow-up examination. The study was approved by the Institutional Ethics Committee of The First Affiliated Hospital of Anhui Medical University with IRB # PJ 2025-11-75. Informed consent for tissue analysis was obtained before surgery. All research was performed in compliance with government policies and the Helsinki declaration.

Paired tumor specimens from a patient with colorectal cancer before and after QLC1101 treatment were obtained from an ongoing phase Ib/II clinical trial conducted at The First Affiliated Hospital of Anhui Medical University (Hefei, Anhui, China). The study was approved by the Institutional Ethics Committee of The First Affiliated Hospital of Anhui Medical University (IRB # PJ 2025-07-13). Tumor responses were evaluated according to the Response Evaluation Criteria in Solid Tumors (RECIST), version 1.1, based on radiographic assessment. Depth of response was defined as the relative change in the sum of the longest diameters of target lesions. Maximal tumor shrinkage after QLC1101 treatment was calculated by comparing the greatest reduction in tumor size with the sum of target lesion diameters at treatment initiation. Written informed consent was obtained from the participant for the collection of clinical data and for publication of this manuscript and any accompanying images.

### Patient derived organoids generation and culture

A total of 16 specimens of human PDAC bearing *KRAS*^G12D^ mutation were acquired from patients who were diagnosed with PDAC between July 2023 and September 2025 at The First Affiliated Hospital of Anhui Medical University. All patients received standard surgical resection and did not receive chemotherapy before surgery. Paraffin-embedded tissues were processed by a pathologist after surgical resection and confirmed as PDAC before further investigation. The study was approved by the Institutional Ethics Committee of The First Affiliated Hospital of Anhui Medical University with IRB # PJ 2025-11-75. Informed consent for tissue analysis was obtained before surgery. All research was performed in compliance with government policies and the Helsinki declaration.

Following pathological confirmation, fresh tumor tissue was processed immediately. Tissue was washed at least three times with ≥10 volumes of wash solution (Advanced DMEM supplemented with 2% FBS, 2 mM EDTA and 10.5 μM Y-27632), minced into fragments <1 mm, and digested in Advanced DMEM containing 5 mg/mL collagenase II, 1 mg/mL dispase and 0.1 mg/mL DNase I at 37 °C with agitation (120 rpm) for 1 h. Digested tissue was filtered through a 70-μm strainer and centrifuged at 300 × g for 5 min. The pellet was washed at least three times, resuspended in wash solution, filtered through a 40-μm strainer, and centrifuged again.

Cells were embedded in Matrigel (D1Med), plated as 50 μL droplets in 24- or 48-well plates, polymerized at 37 °C for 30 min, and overlaid with NGC Organoid^®^ Human Cancer Organoid Culture Medium (pancreatic cancer; D1Med). Medium was refreshed every 2-3 days. Organoids were passaged at 300-500 μm diameter by dissolving Matrigel with ice-cold PBS containing DNase I, followed by digestion with TrypLE at 37 °C for 10-15 min to generate single-cell suspensions for replating or cryopreservation. For cryopreservation, cells were stored in Organoid Cryopreservation Solution (D1Med) in liquid nitrogen. Thawed organoids were recovered using the same dissociation procedure.

### KRAS genotyping of human organoids by Sanger sequencing

During the first passage of organoids, aliquots of cells were reserved for KRAS genotyping. Genomic DNA was extracted using the QIAamp DNA Mini Kit (QIAGEN), followed by PCR-based mutation analysis of KRAS codon 12. Primer sequences were designed as follows: F: 5’-CTGGTGGAGTATTTGATAGTG-3’; R: 5’-CTGTATCAA AGAATGGTCCTG-3’. Sanger sequencing was performed by Sangon Biotech.

### Cell and organoids viability and drug synergy assays

Cells were plated in 96-well plates at a density of 3,000 cells per well and incubated overnight before treatment. Organoids were seeded in 96-well plates at optimal density. Drugs at designated concentrations were added and incubated for 72 h (cell) or 7 days (organoids). Cell viability was determined with a Cell Counting Kit-8 assay (Absin). Optical density was read at 450 nm and analyzed using a microplate reader with SkanIt RE 7.0.1 software (MULTISKAN FC, Thermo Scientific). Organoid viability was determined with CellTiter-Glo assay. Luminescence was analyzed using a microplate reader with SkanIt RE 7.0.1 software. A log[inhibitor] versus response-variable slope model was used to calculate the IC_50_. For all experiments, the medium was replaced every 24 h.

To determine the synergism of two different compounds using viability assays, cells/organoids were plated in 384-well plates at a density of 500 cells (KS and AR of AsPC-1 and No.6) per well and incubated overnight before treatment. Cells/organoids were treated with the indicated combinations of the drugs for 72 h (KS and AR of AsPC-1) or 7 days (KS and AR of No.6) before CCK-8 or CellTiter-Glo assay as manufacturer’s protocol. These experiments were done with four biological replicates. The data were then expressed as percentage inhibition relative to baseline, and the presence of synergy was determined by the Bliss method using the SynergyFinder^+^ web application.

### Immunohistochemistry

Pancreata or livers were surgically removed, fixed in 4% paraformaldehyde in PBS and embedded in paraffin. Five-micrometer sections were prepared and stained with H&E. IHC was performed as before^40^. Slides were scanned using a Motic EasyScanner with MoticEasyScanner software (MOTIC, China). Antibody information is shown in Supplementary Table 1.

IHC scoring was performed as before^40^. Negative and weak staining was viewed as a low expression level and intermediate and strong staining was viewed as a high expression level. For cases with tumors with two satisfactory cores, the results were averaged; for cases with tumors with one poor-quality core, results were based on the interpretable core. Based on this evaluation system, a chi-squared test was used to estimate the association between the staining intensities of ADGRB1-PI3Kγ-pPAK1-CPT1 signaling proteins.

### Luminescence-based ATP detection assay

Cells were seeded in 96-well plates at a density of 3,000 cells per well and allowed to adhere overnight before treatment. Drugs at the indicated concentrations were added for 48 h. Cell number was then determined, and intracellular ATP levels were measured using a luminescence-based ATP detection assay (Promega) according to the manufacturer’s instructions. Luminescence signals were normalized to cell number.

### Serum response factor response element (SRF-RE) luciferase reporter assay

Cells were seeded in 24-well plates and co-transfected with the SRF-RE firefly luciferase reporter plasmid [pGL4.34(luc2P SRF-RE Hygro)] and the pRL-TK renilla luciferase control plasmid using Lipofectamine 3000. Twenty-four hours post-transfection, cells were treated as indicated. Cells were then lysed, and both firefly and renilla luciferase activities were measured using the Dual-Luciferase Reporter Assay System (Yeason) according to the manufacturer’s protocol. Relative luciferase activity was determined by normalizing firefly luciferase signals to renilla luciferase signals.

### Cell and organoid imaging

Cells were cultured on coverslips and fixed in 4% PFA for 10 min at room temperature or methanol for 10 min at −20 °C. Cell immunostaining and macropinosome visualization were performed as before^39^. For lipid droplet (LD) visualization, 2 µM BODIPY (ThermoFisher) or BODIPY-PA (ThermoFisher) together with or without 1 mg/ml high-molecular-mass (70 KDa) TMR-DEX (ThermoFisher) were added to serum-free medium for 30 min (BODIPY) or 60 min (BODIPY-PA) at 37 °C. At the end of the incubation period, the cells were rinsed five times in cold PBS and immediately fixed in 4% PFA. For organoid immunostaining, organoids were fixed directly in 4% PFA for 30 min at room temperature. Each organoid dome was resuspended in 1 mL ice-cold 2% FBS/PBS to remove residual Matrigel, followed by centrifugation at 50 × g for 5 min. This washing step was repeated three times. Organoids were then blocked in 10% FBS/PBS containing 0.1% saponin for 1 h and incubated with primary antibodies overnight at 4 °C on a rocking platform. After washing three times with 2% FBS/PBS and centrifugation at 50 × g for 5 min, organoids were incubated with secondary antibodies and DAPI in 10% FBS/PBS containing 0.1% saponin for 1 h at room temperature with gentle rocking. Organoids were subsequently washed three times with 2% FBS/PBS, centrifuged at 50 × g for 5 min, resuspended in 40 μL antifade mounting medium, mounted onto glass slides and sealed with coverslips. Images were captured and analyzed using a TCS SPE Leica confocal microscope with Leica Application Suite AF 2.6.0.7266 software and analyzed using Huygens and ImageJ. Antibody information is provided in Supplementary Table 1.

### Quantitative PCR analysis

Total RNA was extracted using the Super FastPure RNA Isolation Kit (Vazyme). Reverse transcription was performed using the ABScript II cDNA First-Strand Synthesis Kit (ABclonal). Quantitative (q) PCR was performed as described^40^. Primers were designed using NIH Primer-BLAST (https://www.ncbi.nlm.nih.gov/tools/primer-blast/) and are listed in Supplementary Table 2.

### Western blotting

Tissues, organoids and cells were lysed in RIPA buffer, and protein concentrations were determined using a BCA protein assay kit. Proteins were separated by SDS-PAGE and transferred onto polyvinylidene difluoride (PVDF) membranes. Membranes were incubated with primary antibodies overnight at 4 °C, followed by incubation with horseradish peroxidase (HRP)-conjugated secondary antibodies for 1 h at room temperature. Protein bands were visualized using an enhanced chemiluminescence (ECL) detection system (Tanon) and quantified using ImageJ. Antibody information is provided in Supplementary Table 1.

### RNA-seq library preparation and analysis

Total RNA was isolated from the indicated KS and KR/AR cells. 500 ng of total RNA were enriched for poly-A-tailed transcripts by double incubation with Oligo d(T) Magnetic Beads (NEB) and fragmented for 9 min at 94 °C in 2× Superscript III first-strand buffer containing 10 mM DTT (ThermoFisher). Reverse transcription was performed at 25 °C for 10 min followed by 50 °C for 50 min, and the resulting cDNA was purified using RNAClean XP (Beckman Coulter). Libraries were prepared using either dual unique dual index (UDI) adapters (IDT) or single UDI adapters (Bioo Scientific), PCR-amplified for 11-13 cycles, size-selected using one-sided 0.8× AMPure beads, quantified with the Qubit dsDNA HS Assay Kit (ThermoFisher), and sequenced on a HiSeq 4000 or NextSeq 500 (Illumina) by BGI Genomics.

RNA-seq reads were aligned to the human genome (GRCh38/hg38) using HISAT. Biological and technical replicates were included in all experiments. Transcript quantification was performed using SOAPnuke, and principal component analysis (PCA) was based on transcripts per kilobase million (TPM) values across all genes and samples. Transcript expression values, differential expression analysis, and pathway analyses were performed using Dr. Tom (BGI).

### Fatty acid CoA detection

Indicated KS and KR/AR cells were plated on 10-cm dishes and treated as indicated at ∼80% confluency. Cells were trypsinized, counted, and resuspended in 1 mL ice-cold PBS. 5 mL of ice-cold extraction solvent (methanol:AMBIC:H_2_O, 6:1:3, v/v/v) were added, samples were centrifuged at 2,500 × g for 5 min at 4 °C, and pellets were snap-frozen and stored at −80 °C. Frozen samples were resuspended in 600 μL methanol, homogenized with PTFE beads at 60 Hz for 120 s, mixed with 600 μL chloroform and 240 μL water, incubated on ice for 10 min, and centrifuged at 12,000 rpm for 2 min at 4 °C. Supernatants were dried, reconstituted in 50 μL methanol/water (3:2, v/v), and 30 μL was used for LC-MS analysis.

Separation was performed on a Thermo Scientific Vanquish Core HPLC with a Waters CSH C18 column (100 × 2.1 mm, 1.7 μm) using 10 mM ammonium acetate in water (A) and acetonitrile (B) as mobile phases. Column and autosampler temperatures were 40 °C and 15 °C, respectively, and the injection volume was 2 μL. Ion source settings were: spray voltage 3,200 V, capillary temperature 320 °C, sheath gas 30 arb, auxiliary gas 10 arb, heater 400 °C. Data were acquired with Thermo Xcalibur and quantified using Qual Browser. Sample concentrations were calculated as Cc [ng/mL] × 0.05 mL / 0.43 × cell number. Analyses were performed by the Shanghai Biotree Biomedical Technology.

### Metabolic flow analysis

Indicated KS and KR/AR cells were plated on 10-cm dishes and treated as indicated with 100 μM [U-¹³C]-palmitic acid for 12 h at ∼80% confluency^41^. Cells were harvested by adding 1 mL ice-cold extraction buffer (acetone:methanol:H_2_O, 40:40:20, v/v/v) and scraping. Extracts were transferred to 1.5-mL tubes, stored at −80 °C overnight, and cleared by centrifugation at 15,000 × g for 10 min. Supernatants were used for LC-MS analysis.

Isotope labeling was analyzed using a Shimadzu LC system coupled to a TripleTOF 6600+ mass spectrometer (AB Sciex). Separation was performed on an IHILIC-(P) Classic HILIC column (150 × 2.1 mm, 5 μm, 200 Å) with mobile phase A (95:5 water:acetonitrile containing 20 mM ammonium acetate, 0.1% ammonium hydroxide, 2.5 μM medronic acid) and mobile phase B (acetonitrile). The gradient was: 0-2 min, 85% B; 2-7 min, 85 → 65% B; 7-12 min, 65 → 35% B; 12.1-15.9 min, 20% B; 16-23 min, 85% B; flow rate 0.2 mL/min; injection volume 5 μL; total run time 23 min. MS detection used negative-mode electrospray ionization with GS1/GS2 60 psi, curtain gas 35 psi, ion source temperature 500 °C, and ion spray voltage −4,500 V. Raw data were processed using El-MAVEN (v0.12.1) to integrate ¹²C and ¹³C isotopologue peak areas, corrected for natural isotope abundance using AccuCor (R package). Labeling fractions were calculated as, Labeling fraction = [Area of specific isotopologue] / Σ[Areas of all isotopologues]. Analyses were performed by the Institute of Metabolism & Integrative Biology, Fudan University.

### Air flow-assisted desorption electrospray ionization mass spectrometry imaging (AFA-DESI-MSI) assay

Orthotopic PDAC mouse models were generated as described above. From day 10 post-transplantation, mice were treated with DOX (1 mg/mL, drinking water) or DOX and MRTX for 4 days and euthanized 4 h after the final dose. Tumor tissues were harvested, snap-frozen in liquid nitrogen vapor, and sectioned into 10 μm cryosections, which were mounted onto Superfrost Plus slides (Thermo Fisher Scientific). Prior to analysis, sections were stored at −40 °C overnight and dehydrated under vacuum.

AFADESI-MSI was performed using an air flow-assisted desorption electrospray ionization platform (AFAI-MSI, Viktor) coupled to a Q Exactive Orbitrap HFX mass spectrometer (Thermo Fisher Scientific). The electrospray solvent (acetonitrile/water, 90:10, v/v) was delivered at 3 μL/min. Data were acquired in positive and negative ion modes (±4.5 kV) using nitrogen as the spray gas (0.65 MPa). The spray angle was 60°, with distances of 6 mm from sprayer to tissue and 4 mm from sprayer to inlet capillary; the capillary temperature was maintained at 350 °C.

Full-scan spectra were collected over an m/z range of 70-1050 at a resolution of 60,000. The sample stage was rastered at 0.25 mm/s with a step size of 50 μm, defining the spatial resolution. Data acquisition was controlled using Xcalibur software (Thermo Fisher Scientific). AFADESI-MSI analyses were performed by Shanghai Majorbio Bio-Pharm Technology.

### Matrix-assisted laser desorption/ionization mass spectrometry imaging (MALDI MSI) assay

Orthotopic PDAC mouse models were established as described above. From day 10 post-cell transplantation, mice were treated with DOX (1 mg/mL, drinking water) or DOX and MRTX for 4 days and euthanized 4 h after the final dose. Tumors were harvested, cut into 350 μm slices using a vibratome (Leica), and incubated in 10% lipid depleted FBS/DMEM containing 100 μM [U-¹³C]-palmitic acid for 4 h^42,43^. Treated slices were snap-frozen and sectioned into 10 μm cryosections. Sections for MALDI MSI were thaw-mounted on indium tin oxide (ITO)-coated glass slides (Bruker) and desiccated under vacuum for 10 min. Alternating sections were collected on standard slides for IHC or immunofluorescence.

Tissue sections on ITO slides were sprayed with 10 mg/mL NEDC in 70:30 methanol:water using an HTX TM sprayer (HTX Technologies) at 80 °C, 0.1 mL/min flow rate, 1,000 mm/min velocity, 2 mm track spacing, 10 psi pressure, and 3 L/min gas flow. Ten passes were applied with 10 s drying between passes. MALDI-FT-ICR MS imaging was performed on a solariX XR FT-ICR mass spectrometer (9.4 T, Bruker Daltonics) with resolving power of 120,000 at m/z 500. Mass calibration was performed with 1 mg/mL arginine to ≤1 ppm accuracy, and the x-y raster width was set to 50 μm. Data acquisition and analysis were performed using mMASS, FlexImaging, and SCiLS Lab (Bruker Daltonics). Analyses were performed by the Institutional Center for Shared Technologies and Facilities of Institute of Nutrition and Health, Chinese Academy of Sciences.

### Seahorse assay

Cells were plated on 96-well and XF96-well plates at 5,000 cells/well in DMEM containing 10% FBS and cultured overnight for adherence. For the fatty acid oxidation stress test, cells were starved in substrate-limited XF DMEM (0.5 mM glucose, 1 mM glutamine, 0.5 mM L-carnitine, 1% FBS) with indicated drugs for 12-16 h. Sensor cartridges were hydrated overnight in XF Calibrant (Agilent) at 37 °C in a non-CO_2_ incubator and loaded with etomoxir (40 μM), oligomycin (15 μM), FCCP (15 μM), and rotenone/antimycin A (5 μM) in ports A-D. Cells were incubated in assay medium (0.5 mM L-carnitine, 2 mM glucose, 100 μM palmitic acid) and transferred to the XFe96 analyzer for OCR measurement. For FAO assay, cells were incubated in assay medium (0.5 mM L-carnitine, 2 mM glucose), and cartridge ports were loaded with Veh/palmitic acid (100 μM), oligomycin, FCCP and rotenone/antimycin A in ports A-D. For mitochondrial stress test, cells were cultured in complete medium, and assay medium (XF DMEM with 10 mM glucose, 1 mM pyruvate, 2 mM glutamine) was used; cartridge ports were loaded with oligomycin, FCCP and rotenone/antimycin A in ports A-C. OCR was calculated using Seahorse Wave Controller software (v2.6.3.5, Agilent) and normalized to cell number using CCK-8 or BCA assays.

### Lipid uptake assays

Fatty-acid transport kinetics were assessed as previously described^44^. Indicated KS and KR/AR cells were seeded in 96-well plates at 5,000 cells/well and cultured for 2 days to 95-100% confluence. Cells were serum-starved in DMEM for 1 h before the assay. Medium was replaced with complete medium containing BODIPY-palmitic acid. Extracellular fluorescence was quenched with Trypan blue to eliminate non-cell-associated signals. Fluorescence kinetics (Ex/Em = 485/528 nm) were recorded every 50 s for 1 h using a Thermo Fisher microplate reader with SkanIt RE 7.0.1 software. Data were normalized to the initial fluorescence at time 0.

### Lipid transfer experiments

3T3-L1 cells were differentiated and labeled with 2 μM BODIPY-palmitic acid for 4 h^45^. For *in vitro* lipid transfer assays, differentiated 3T3-L1 adipocytes were cultured on the upper chamber of a Transwell system (Labselected), while indicated KS and KR cells were plated on 12-mm circular coverslips in the lower chamber and allowed to adhere. After washing, BODIPY-labeled adipocytes were transferred to the lower chambers containing KS or AR cells, and co-cultures were treated as indicated for 24 h. KS or AR cells were then fixed, stained, mounted, and imaged using a Leica TCS SPE confocal microscope with Leica Application Suite AF software. Images were analyzed using Huygens and ImageJ.

For *in vivo* lipid transfer assays, orthotopic tumor models were established as described above. On day 7 post-cell transplantation, mice were intraperitoneally injected with 1 × 10^7^ BODIPY-palmitic acid-labeled 3T3-L1 adipocytes. After 24 h, tumors were harvested, digested in DMEM containing 5 mg/mL collagenase II, 1 mg/mL dispase, and 0.1 mg/mL DNase I, stained with anti-EpCAM antibody, and analyzed by flow cytometry using an Attune X cytometer (Thermo Scientific). Data were analyzed using FlowJo.

### Statistics and reproducibility

Macropinosomes and mitochondria were quantified using the ‘Analyze Particles’ feature in ImageJ (NIH). The macropinocytotic uptake index^46^, mitochondrial and LD content, as well as mitochondria-LD and macropinosome-LD co-localization, were calculated by normalizing the specific organelle area to the total cell area per field. Data represent the average across 15-20 fields. Tumor area (%) was quantified by using the ‘Polygon’ and ‘Measure’ feature in Fiji Image J and was computed by tumor area in relation to total area for each field and then by determining the average across all the fields (6-7 fields). Protein-positive area (%) was quantified in Fiji (ImageJ) using the Color Deconvolution (H DAB) and Analyze Particles functions and calculated as the ratio of protein-positive area to total tissue area per field, followed by averaging across 5-7 randomly selected fields. Organoid growth was quantified by measuring the projected area of individual organoids using the Polygon and Measure tools and normalized to the area of the same organoid on day 1 (10 organoids per condition). Relative metabolite abundance was quantified using adjacent H&E-stained sections to identify tumor regions. For each field, signals were normalized to total ion current (TIC), and values were averaged across six fields. All measurements were performed in randomly selected fields of view.

Statistical analyses were performed using GraphPad Prism. Two-sided t-tests, Mann-Whitney tests, one-way ANOVA, Kruskal-Wallis tests, or Brown-Forsythe and Welch ANOVA were used as indicated. Data are presented as mean ± s.e.m. Kaplan-Meier survival curves were analyzed using the log-rank test. Correlations between ADGRB1-PI3Kγ-pPAK1 signaling proteins in human PDAC specimens were assessed using two-tailed chi-squared tests. Statistical significance was defined as \*\*\*\**P* < 0.0001, \*\*\**P* < 0.001, \*\**P* < 0.01, and \**P* < 0.05. All experiments, except immunohistochemical analysis of human specimens, were repeated at least three times.

### Data availability

RNA-seq data are available at the Genome Sequence Archive for Human (GSA-Human) under accession number (HRA016532). Graph data and raw images of immunoblot are provided within the Source Data. All raw image data including immunostaining, immunoblotting, IHC, and H&E staining were uploaded to Mendeley Data (10.17632/3dfmh6847y.1). Source data are provided with this paper.

